# Interplay of sleep neural oscillations enhances coordinated memory reactivation between cortex and hippocampus

**DOI:** 10.64898/2026.06.12.731367

**Authors:** Masahiro Takigawa, Diao Tong, Edward A.B. Horrocks, Aman B. Saleem, Daniel Bendor

## Abstract

Memory consolidation during sleep requires coordinated reactivation of specific experiences across hippocampus and cortex. This process occurs with synchronized neural oscillations including cortical slow waves, thalamocortical spindles, and hippocampal sharp-wave ripples. While temporal coupling of these rhythms is implicated in consolidation, a fundamental question remains: do oscillations reflect a general increase in communication, or do they selectively coordinate which memories are reactivated together across regions? Here we segregated competing memory representations in primary visual cortex by training mice on two visually distinct virtual reality tracks, each restricted to one visual hemifield, rendering their representations lateralized in cortex. Using large-scale electrophysiology, we demonstrate that hippocampus and cortex coherently reactivate the same memories. Temporally, the cortical memory trace active before and after a ripple is coherent with the memory hippocampus reactivates. Crucially, this reactivation coherence is maximally enhanced by high hippocampal ripple power in concert with high cortical spindle-band power and slow oscillation trough phase localized to the dominant memory trace reactivating. Our findings establish that sleep oscillations coordinate content-specific cortico-hippocampal reactivation necessary for consolidation.

## Introduction

During sleep, the hippocampus and cortex require bidirectional communication to consolidate recent labile memories into stable long-term representations^1–10^, a mechanism that must operate at a scale sufficient to support multiple episodic events and contexts from waking experience^11^. To consolidate during sleep, communication between hippocampus and cortex should be coordinated and selective, ensuring that the same memory trace is reactivated across both regions. Memory consolidation during sleep is thought to occur mainly during the non-rapid-eye-movement (NREM) stage, which is dominated by synchronized brain activity, manifesting in three stereotyped oscillations: cortical slow-oscillations^12^ (SOs, 0.5-4Hz), thalamocortical spindles^13^ (9-17Hz), and hippocampal sharp-wave ripples^14^ (SWRs, 125-300Hz), whose temporal coupling is postulated to enhance the cortico-hippocampal communication required for consolidation^9,15–17^. However, how these oscillatory dynamics support the selective reactivation of specific memories across both cortex and hippocampus is not known. We hypothesize that the interplay of ripples, local spindles, and slow oscillations provides a window of spatial and temporal engagement, allowing coordinated reactivation of the same memory across brain regions.

Demonstrating this interaction has, so far, been challenging as studying selective reactivation requires two or more distinct experiences^7,18–20^, which are typically encoded by overlapping neuronal populations within the same cortical circuits. As a result, it is challenging to disentangle whether the influence of a local cortical oscillation is directed toward the reactivation of a specific memory trace or simply provides a broader, non-selective modulation of cortical activity. To overcome these challenges, we physically segregated two competing memory traces in cortex, so any given local cortical oscillation is strongly biased to interact with one trace over the other. We achieved this by two visually distinct virtual reality tracks, each restricted to half of the mouse’s visual field, thereby rendering each track-specific representation hemispherically biased in primary visual cortex (V1). Using large-scale electrophysiology, this design allowed us to ask how ongoing oscillatory dynamics influence the coherence of reactivation content between V1 and hippocampus during sleep.

We discover that hippocampus and cortex coherently reactivate the same content (i.e. memory trace of the specific track). This reactivation coherence was maximized by of hippocampal ripple power combined with local cortical spindle-band power and slow-oscillation phase in the hemisphere representing the reactivated memory. Temporally, we found that cortical content both before and after ripple onset predicts hippocampal reactivation content. Crucially, each of these three oscillatory features makes a distinct, non-redundant contribution to cortico-hippocampal reactivation coherence, collectively defining a spatio-temporal window that enables a content-specific communication during consolidation.

## Results

### Creating separable, context-specific cortico-hippocampal representations

To create two physically segregated memory traces in visual cortex, we designed two closed-loop virtual reality (VR) linear tracks that head-fixed mice (n=6) navigated (Fig. 1a). Crucially, we designed the linear tracks to be visually distinct by having them cover different visual hemifields – each VR track only had visual cues on one visual hemifield, with the opposite hemifield being gray (Track L-visual cues left, Track R-visual cues right) (Fig. 1a). We simultaneously recorded bilaterally from primary visual cortices (V1) and CA1 region of the hippocampus using chronically implanted 4-shank Neuropixels 2.0 probes21,22 (Extended Data Fig. 1) during VR exploration and unrestrained sleep after that experience. Confining each track to one visual hemifield rendered V1 representations highly separable, with each track predominantly lateralized to the contralateral hemisphere. Neurons from left V1 (total n = 1539 cells; mean ± SE = 70.0 ± 8.2 per session) preferentially responded to the right-sided track (i.e. Track R) while neurons from right V1 (total n = 1617 cells; mean ± SE = 73.5 ± 6.2 per session) preferentially responded to the left-sided track (i.e. Track L) (Fig. 1b, Extended Data Fig. 2). Hippocampal neurons (total n = 1564, mean ± SE = 71.1 ± 6.4 per session) also represented both tracks, but unlike V1, there was no obvious hemispheric bias for track preference (Fig. 1b, Extended Data Fig. 2). The track the animal was currently experiencing could be decoded using either V1 or hippocampal activity, based on a population decoder (Fig. 1c,d; see Methods for decoder details; Extended data Fig. 3;V1 AUC: real vs. track-label shuffle 95^th^ percentile, mean ± SE = 0.957 ± 0.0115 vs. 0.515 ± 5.64×10^−4^; Wilcoxon signed-rank test, p = 2.15×10^−5^; hippocampal AUC: real vs. shuffle, mean ± SE = 0.870 ± 0.0178 vs. 0.515 ± 5.92×10^−4^; Wilcoxon signed-rank test, p = 2.15×10^−5^).

**Fig 1.**
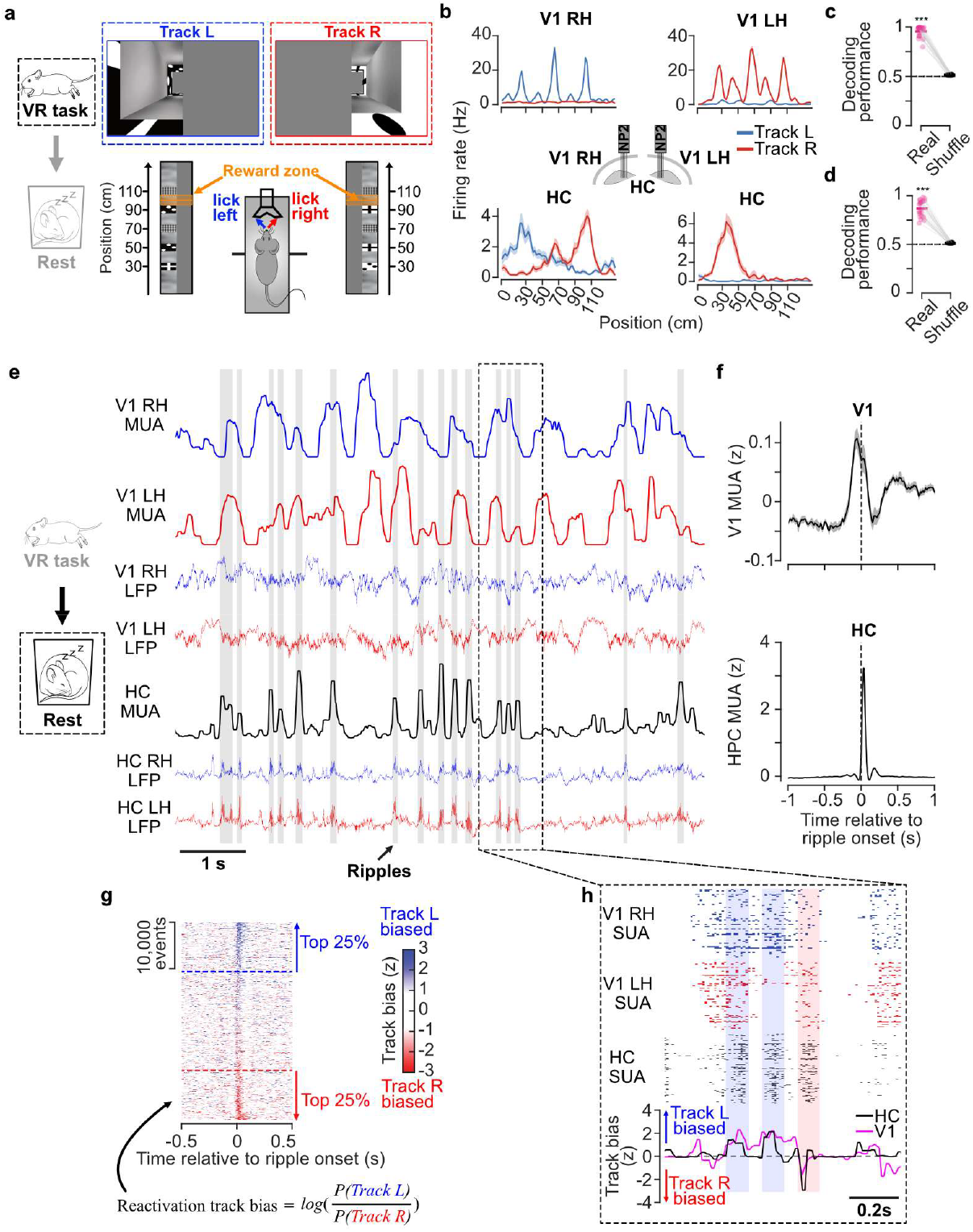
Creating separable, context-specific cortico-hippocampal representations. **a**.Schematic of the two-track virtual reality (VR) paradigm. Head-fixed mice navigated two VR linear tracks by running on the wheel for a reward in a fixed position. Visual features in each track were designed to occupy half of the screen (with the other half grey) - screenshots show visual scenes at position 30cm for both Track L (top left) and Track R (top right). Animals licked left and right to receive liquid rewards in a reward zone (95-105cm). **b**.Spatial tuning curves of example neurons from V1 left hemisphere (V1 LH, top left), V1 right hemisphere (V1 RH, top right) and hippocampus (HC, bottom two examples) for both Track L (blue) and Track R (red). Note that V1 represents the contralateral visual field. **c-d**. Track identity decoding performance based on decoded activity from either **(c)** V1 or **(d)** HC spiking using area under the receiver operating characteristic curve (ROC AUC) with 10-fold cross-validation. Observed data (pink) are compared against track-label shuffled data (grey; 95th percentile). Each point represents one session; horizontal lines indicate the mean across sessions. ***p = 2.15×10^−5^ for V1 and HC AUC real vs. track-label shuffle, two-tailed Wilcoxon signed-rank test. **e**. Example local field potential (LFP) and neuronal spiking activity from bilateral V1 and HC during sleep after the experience. LFP and spikes from left and right hemispheres are shown in red and blue, respectively. Bilateral HC spikes were shown in black. Detected sharp-wave ripples (SWRs) are shaded in grey. **f**. Mean multi-unit activity (z-scored) from V1 (top) and HC (bottom) relative to ripple onset across sessions (n = 22 sessions). The shaded regions indicate the standard error across sessions. **g**. Schematic representation of the main research questions: How does ripples, spindles and slow-oscillations influence the coherence of context-specific reactivation between V1 and HC?**h**. Distribution of hippocampal reactivation bias during ripple events. Reactivation track bias for each time bin is calculated as the log odds of the decoded probability of Track L *(+ve*, more blue) over Track R (*-ve*, more red) (see method). Track bias is plotted as a function of time from ripple onset for all ripple events (n = 42,466 ripple events from 22 sessions). Events with the top 25% of positive (blue) and negative (red) HC bias are selected as Track L– and Track R–biased ripple events, respectively. **i**. Spiking activity and track bias of V1 and HC from example reactivation events. Example Track L biased (shaded in blue) and Track R biased (shaded in red) ripple events are labelled based on HC track bias. For **a** and **e**, mouse schematics were adapted from SciDraw.io (http://doi.org/10.5281/zenodo.3925949, http://doi.org/10.5281/zenodo.3926215).

Following experience on the two VR tracks, we recorded V1 and hippocampus activity from unrestrained mice in a sleep box. Consistent with previous work^5,6^, during sleep, we observed spindles and slow oscillations in V1 where the neuronal population had alternating epochs of higher and lower firing rates. In hippocampus, we observed sharp-wave ripples (total = 42,466 ripple events, mean±SE =1930 ± 136.1 events per session; see method; Extended data Fig. 4), accompanied by transient increase in the population spiking activity (Fig. 1e,f). Both hippocampal and V1 overall spiking activity were generally increased peri-ripple onset (Fig. 1f). While hippocampal activity was tightly restricted to the ripple window, V1 activity was elevated over a wider time window, generally increased 200 ms prior to ripple onset, and persisting beyond ripple onset (about 100 to 200 ms after ripple onset). We therefore questioned whether the content of V1 activity (i.e. memory traces of the specific track) before and after ripple onset coherent with the HC reactivation content during ripple?

To quantify whether the content of reactivations in V1 and hippocampus were matched, we applied a probabilistic decoding approach to independently calculate reactivation likelihood for each track in V1 and hippocampus. To measure which track was currently represented in the neural population, we calculated track bias using a log-odds metric^5–7^, which represented track identity and likelihood by its sign and magnitude, respectively (e.g. large positive value-high likelihood of Track L, small negative value-low likelihood of Track R). This track bias decoding metric allowed us to identify track-biased ripple events based on the mean hippocampal (HC) track bias during each individual ripple, where the positive values corresponded to Track L bias and the negative values to Track R bias (Fig. 1g). Using this approach, we observed many instances of coherent V1-HC re-activation around the time of ripple onset, where the direction and magnitude of track bias in hippocampus correlate with those in V1 (Fig. 1i; Extended Data Fig. 5)

We first establish that the content of reactivation events can be coherent between hippocampus and cortex, and next investigate how local cortical oscillations influence this coherence between V1 and hippocampus.

### Hippocampus and cortex reactivate the same content, especially during events with high ripple power

While V1 activity is coupled to hippocampal ripples^6,23^, when multiple memories are reactivating, it remains unclear whether hippocampus and V1 reactivate the same memory during ripples. To investigate this, we determined whether the activity of individual V1 neurons depended on which track the hippocampus reactivated. We found that V1 neurons that preferred one track during behavior exhibited higher activity when the same track reactivated in HC. (defined by the upper and lower quartiles of the HC track bias distribution reactivating Track L and Track R, respectively) (Fig. 2a). To quantify this effect, we computed the mean firing rate difference between tracks for each V1 neuron during behaviour and compared it to the difference during ripples classified as Track L or R based on HC decoded track bias (Extended Data Fig.6). Using a linear mixed-effects model, we found that firing-rate differences between the two tracks during running predicted the firing-rate difference during hippocampal reactivation events (-200 to 200ms relative to ripple onset, β = 0.079, F_(1,3145)_ = 27.1, p = 2.10×10^−7^, R^2^ = 0.30; Extended Data Fig.6). This predictive relationship remained significant both for V1 activity prior to (−200 to 0ms relative to ripple onset, β = 0.057, F_(1,3145)_ = 13.6, p = 2.31×10^−4^, R^2^ = 0.28; Extended Data Fig.) and following ripple onset (0 to 200ms relative to ripple onset, post-onset, β = 0.088, F_(1,3145)_ = 32.1, p = 1.56×10^−8^, R^2^ = 0.27; Extended Data Fig.6).

**Figure 2.**
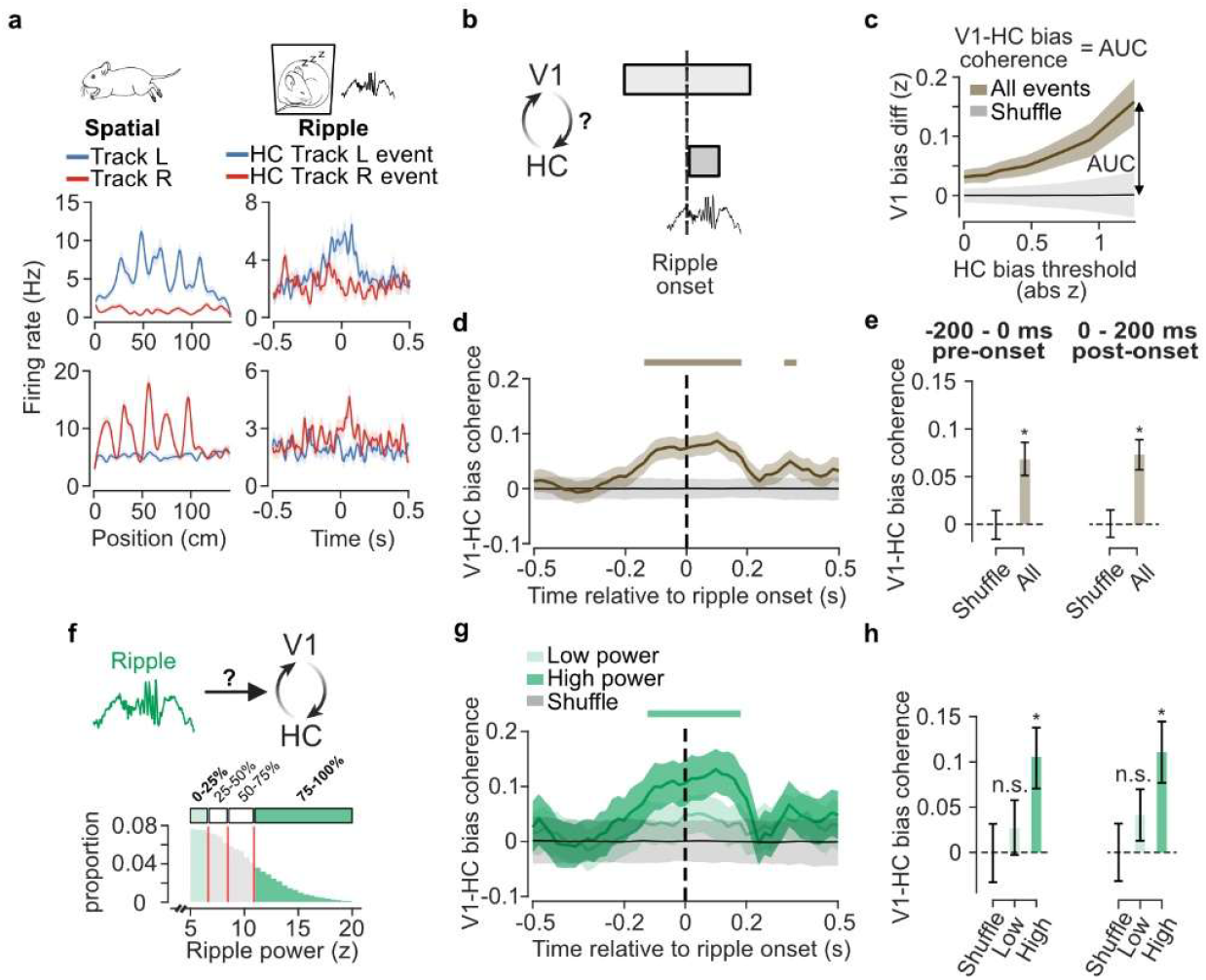
V1 and hippocampus reactivate the same content especially during ripples with high ripple power. **a**. Example spatial tuning (left) for Track L (blue) and Track R (red) and ripple PSTH (right) for Track L (blue) and Track R (red) biased ripple events (defined by top 25% hippocampal track bias) from a Track L preferring (top) and Track R preferring (bottom) V1 neuron. **b**. Schematic of the experimental question: Does hippocampus and V1 reactivate the same content (i.e. track)? **c**. V1 bias difference between HC Track L and Track R biased ripple events at various HC bias thresholds (-0.05 to 0.05s window) for all ripple events (brown) and shuffled controls (gray). Area under the curve (AUC) of the V1 bias difference curve was used to index V1-HC reactivation bias coherence. **d**. Time course of V1-HC reactivation bias coherence relative to ripple onset for all ripple events (100ms sliding window with 20ms steps). The shaded region indicates the 95% bootstrapped confidence interval (CI) of the selected ripple events across all sessions. The shaded region in gray indicates the 95% CI of the event label shuffle. Colored bars above indicate time windows of significance where the 95% CI of the observed data exceeded the 95% CI of the shuffled distribution. **e**. V1-HC bias coherence for the pre-onset window (-200 to 0ms, left) and post-onset window (0 to 200ms, right). **f**. Distribution of ripple power at ripple peak for ripple events across all sessions. Events with low power (lowest quartile) and high power (highest quartile) were highlighted. Reactivation events for later analysis were selected based on HC bias following ripple onset (0–100ms) as well as ripple power. **g**. Time course of V1-HC reactivation bias coherence relative to ripple onset for low (light green) and high (dark green) ripple-power events (100ms sliding window with 20ms steps). The shaded regions indicate the 95% bootstrapped CI of the selected ripple events across all sessions. The shaded region in gray indicates the 95% CI of the event label shuffle. Colored bars above indicate time windows of significance where the 95% CI of the observed data exceeded the 95% CI of the shuffled distribution. **h**. V1-HC bias coherence for the pre-onset window (-200 to 0ms, left) and post-onset window (0 to 200ms, right). Error bars represent the 95% bootstrapped CI of the selected events or the 95% CI shuffled distribution. Asterisks (*) denote significant coherence where the 95% CI of the real data does not overlap with the 95% CI shuffled distribution.

We also sought to investigate if the content of cortico-hippocampal reactivation was coherent at population level. To quantify the reactivation coherence between V1 and hippocampus, we first selected ripple events classified as Track L and Track R using the HC track bias during the ripple (0 - 0.1s), and then calculated the track bias difference for V1 reactivations (i.e. Track L bias minus Track R bias) at different times relative to ripple onset (Fig. 2b). To ensure the results were not dependent on an arbitrary bias threshold for event selection, we systematically shifted the HC bias threshold (10 deciles) for event selection and examined how the mean V1 track bias difference varied with the HC thresholds at different times relative to ripple onset (100ms sliding window, 20ms step) (Fig. 2c). By doing so, we observed the track bias difference in V1 increased as the HC bias threshold for event selection tightened (Fig. 2c, e.g. - 0.05 to 0.05s). Such a relationship was abolished when the same procedure was repeated using event label shuffling (Fig. 2c). Using this approach, we calculated the area under this track bias difference curve (AUC) to infer the V1-HC re-activation coherence, providing a threshold-free metric. We observed that V1-HC coherence was significantly elevated above shuffled baseline both before and after ripple onset (-140 to 180 ms, based on 95% bootstrapped confident interval. Fig. 2d). Taking 200ms window before (-200 to 0ms, pre-onset) and after (0 to 200ms, post-onset) ripple onset, the 95% confidence interval of the V1-HC coherence measure was significantly higher than the event label shuffled distribution (Fig. 2e).

We next examined how the amplitude of hippocampal ripples influenced the context-selective ripple modulation of V1. As high-power ripple events are observed to enhance memory reactivation and memory recall^17^, we asked whether ripple power increased the coherence of the V1-HC reactivation content (Fig. 2f). To investigate this, we divided ripple events into quartiles depending on peak ripple power (Fig. 2f). The reactivation coherence between V1 and hippo-campus was increased by ripple power. We observed that V1-HC coherence was significantly elevated above shuffled baseline both before and after ripple onset for high-power events but not for low-power events (high ripple power: from -120 to 180ms, low ripple power: n.s. Fig. 2g). Taking 200ms window before (-200 to 0ms, pre-onset) and after (0 to 200ms, post-onset) ripple onset, the 95% confidence interval of the V1-HC coherence overlapped with the event label shuffled distribution for low-power events but not for high-power events (Fig. 2h). Similarly, at neuronal level, the predictive relationship for track-selective ripple modulation of V1 neuronal firing rate was significant for the highest power quartiles (-200 to 200ms relative to ripple onset, β = 0.11, F_(1,3145)_ = 46.6, p = 1.04×10^−11^, R^2^ = 0.27; Extended Data Fig.6), and not for the lowest power quartiles (-200 to 200ms relative to ripple onset, β = 0.0025, F_(1,3145)_ = 0.024, p = 0.88, R^2^ = 0.18; Extended Data Fig.6). Together, these findings suggest that hippocampus and cortex reactivated the same memory, especially during events with high ripple power.

### Coherence of cortico-hippocampal reactivation content depends on local spindle-band power

In addition to hippocampal ripple power, how do ongoing cortical oscillations influence the coherence of cortico-hippocampal reactivation? Cortical spindle-band (9–17Hz) oscillations are known to couple with hippocampal ripple activity to support memory consolidation^9,15,17^. Hippocampal ripples are often nested within periods of high cortical spindle-band power^8,10,15,24^ and disrupting coupling of ripples and spindles can lead to memory impairment^15,25^. We therefore examined whether local V1 spindle-band power modulated the coherence of cortico-hippocampal reactivation during ripple events (Fig. 3a). Our paradigm was uniquely suited to this question due to artificially creating a hemispheric bias in reactivation content in our behavioural paradigm: Track R was primarily encoded in the left V1, while Track L was primarily encoded in the right V1. This hemispheric difference allowed us to test whether there was a selective enhancement of HC–V1 coherence by local V1 spindle-band power in the dominant hemisphere, which we define as the V1 hemisphere associated with the reactivation track bias in V1 (i.e. strong V1 bias towards Track L paired with V1 spindle-band power from contralateral right hemisphere). We divided V1 spindle-band power at ripple peak into quartiles (Fig. 3b) and selected Track L and Track R biased events based on V1 track bias, identifying the dominant hemisphere for each event (Fig. 3b,c). To infer V1-HC reactivation coherence, we quantified the HC track bias difference between Track L and Track R biased events, defined by V1 track bias, for the lowest and highest spindle-band power quartiles in the dominant hemisphere (Fig. 3d). The reactivation coherence between V1 and hippocampus increased with the local spindle-band power in the dominant hemisphere. For ripple events with high local spindle-band power, V1-HC coherence was significantly elevated above the shuffled baseline across a wide window spanning both before and after ripple onset (−120 to 180 ms; Fig. 3e). By contrast, low spindle-band power was associated with a markedly narrower window of significant re-activation coherence, restricted to a brief period following ripple onset (0.06 to 0.08s; Fig. 3e). This distinction was further confirmed by examining V1-HC coherence in 200 ms pre-onset (−200 to 0 ms) and post-onset (0 to 200 ms) windows: for high spindle-band power events, reactivation coherence was significantly above the shuffled distribution in both windows, whereas for low spindle-band power events, it failed to exceed the shuffled distribution in either window (Fig. 3f). The results suggested that the reactivation coherence between V1 and hippocampus was higher when ripples were associated with higher local spindle-band power.

**Figure 3.**
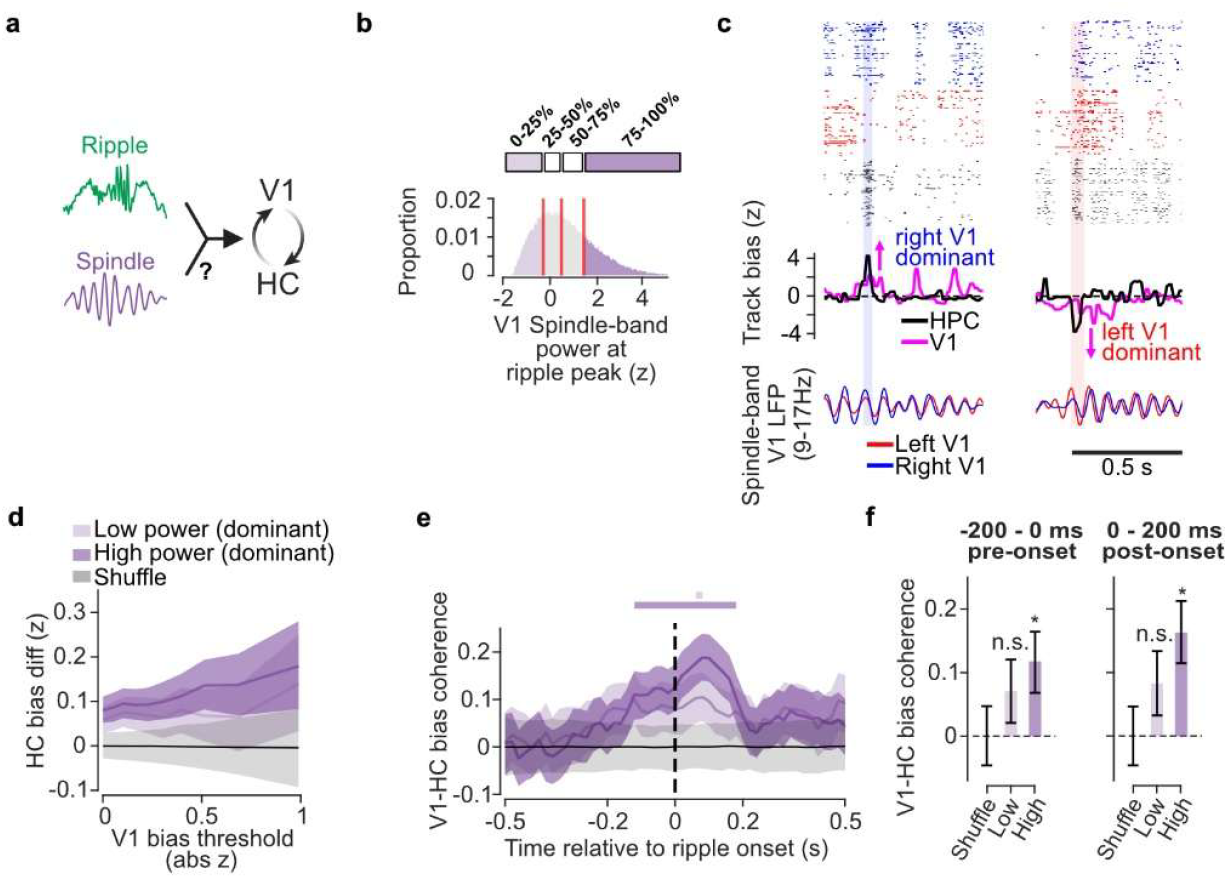
Coherence of cortico-hippocampal reactivation content during ripple depends on the spindle-band power. **a**. Schematic of the experimental question: Does V1 spindle-band power during ripple modulates the coherence of reactivation content between the hippocampus (HC) and V1.**b**. Distribution of V1 spindle-band power at ripple peak. Events with spindle-band power events with low (lowest quartile) and high (highest quartile) were highlighted. Reactivation events for later analysis were selected based on V1 bias as well as the V1 spindle-band power in the dominant hemisphere (i.e. the hemisphere that strongly contributed to the V1 track bias for each event) **c**. Example coherent Track L biased events (shaded in blue, left) with high spindle-band power in right V1 (dominant hemisphere, blue). Example coherent Track R biased events (shaded in red, right) with high spindle-band power in left V1 (dominant hemisphere, red). **d**. HC bias difference between V1 Track L and Track R biased events at various V1 bias thresholds (-0.05 to 0.05s window) for low (light purple) and high (dark purple) power events in context-matching hemisphere. AUC under the HC bias difference curve was used to index V1-HC reactivation bias coherence. **e**. Time course of V1-HC bias coherence relative to ripple onset for low (light purple) and high (dark purple) power events (100ms sliding window with 20ms steps). The shaded regions indicate the 95% bootstrapped confidence interval (CI) of the selected ripple events across all sessions. The shaded region in gray indicates 95% CI distribution of the event label shuffle. Colored bars above indicate time windows of significance where the 95% CI of the observed data exceeded the 95% CI of the shuffled distribution. **f**. V1-HC bias coherence for V1 bias from pre-onset window (-200 to 0ms, left) and post-onset window (0 to 200ms, right). Error bars represent the 95% bootstrapped CI of the selected events or the 95% CI of the event label shuffled data. Asterisks (*) denote significant coherence where the 95% CI of the real data does not overlap with the 95% CI shuffled distribution.

### Cortical activity before ripple predicts hippocampal content when there are local asymmetries in slow-oscillations

In addition to spindle-band oscillations, slow-oscillations (SOs, 0.5–4Hz) are also known to couple with hippocampal ripple activity to support memory consolidation^8–10,26–28^. Given that neuronal populations are more active during the SO trough phase^12,29^, we asked whether the phase of cortical SOs in the dominant hemisphere (i.e. the hemisphere associated with the V1 track bias) influences the coherence of cortico-hippocampal reactivation (Fig. 4a). To investigate this, we divided ripple events based on whether they occurred closer to the SO trough phase (-π to -π/2 or π/2 to π; left: n = 17888; right: n = 14900) or peak phase (-π/2 to π/2; left: n = 24578; right: n = 27566) (Fig. 4b). We then selected events based on V1 track bias and SO phases in the dominant hemisphere (Fig. 4c). Similar to previous reports^12,29^, the V1 activity itself was modulated by V1 SO phase such that the mean firing rate was higher near SO trough phases compared to SO peak phases (Fig. 4d; Ipsilateral peak = 2.38±0.19 Hz, Ipsilateral trough = 4.21±0.32 Hz, p = 4.01×10^−5^, Wilcoxon signed rank test).

**Figure 4.**
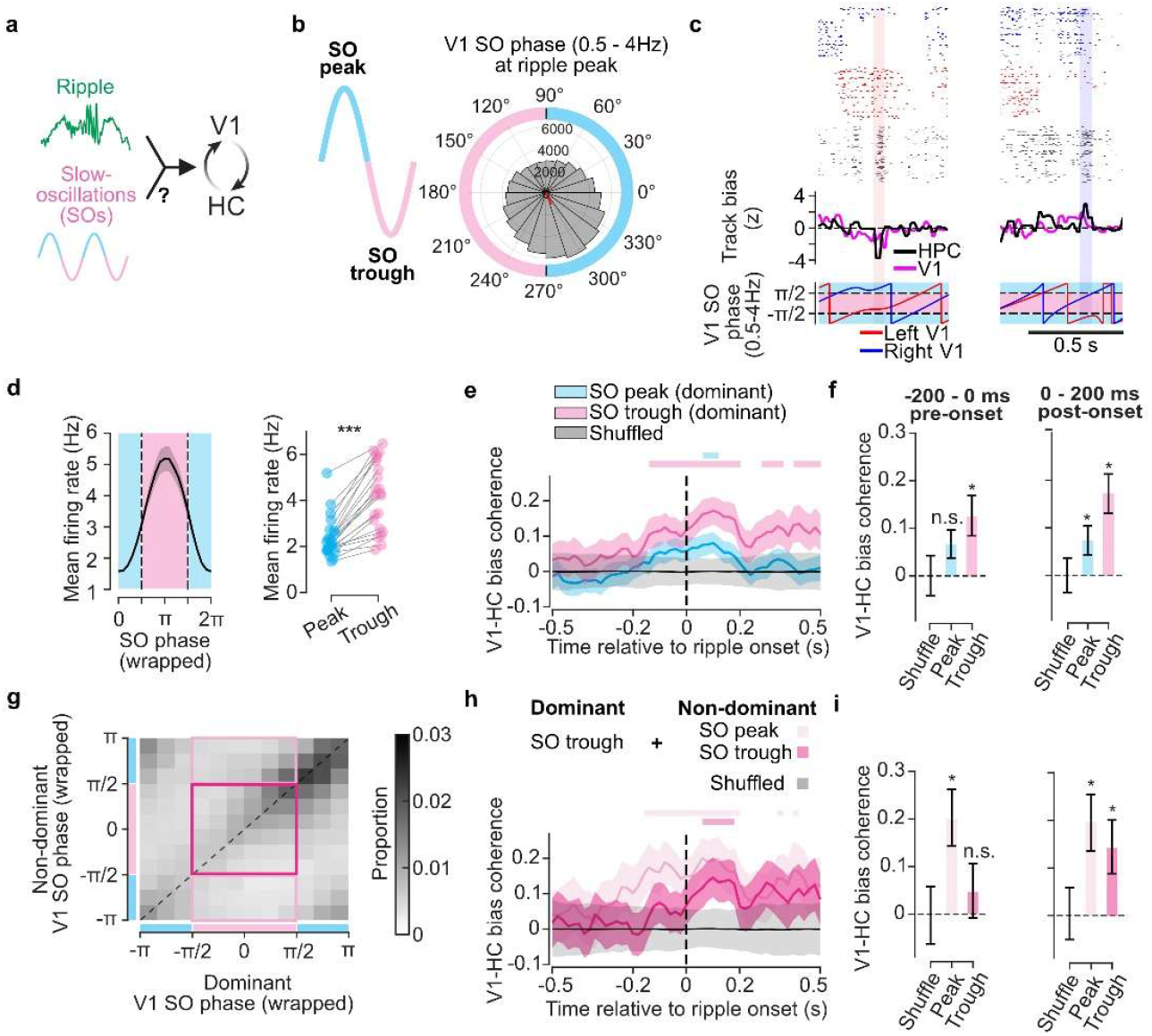
Pre-ripple cortical bias predicts subsequent hippocampal content when there is asymmetry in cortical slow-oscillations. **a**. Schematic of the experimental question: Does V1 SO phase during ripple modulates the coherence of reactivation content between the hippocampus (HC) and V1 **b**. Distribution of V1 SO phase at ripple peak. n = 42466 events, preferred angle = 293°, r = 0.19, p < 10^−300^. Phases near the peak phase (-π/2 to π/2) and the trough phase (-π to -π/2 or π/2 to π) are highlighted in blue and pink. Reactivation events for later analysis were selected based on V1 bias as well as V1 SO phase (0.5-4Hz) at ripple peak. **c**. Example coherent Track L biased events (shaded in blue, left) with SO trough in the right V1 (dominant hemisphere, blue). Example coherent Track R biased events (shaded in red, right) with SO trough in the left V1 (dominant hemisphere, red). SO phases near the peak phase and the trough phase are shaded in sky blue and pink. **d**. Mean V1 firing rate at different SO phases. (Left) Mean firing rate of V1 neurons relative to the ipsilateral V1 SO phases. The line and shaded regions indicate mean and standard error across 22 sessions. (Right) Mean firing rate during the peak (sky blue) vs trough (pink) phase state.***p = 4.01×10^−5^ for ipsilateral V1 peak vs. trough, two-tailed Wilcoxon signed-rank test. **e-f**. V1-HC reactivation bias coherence for events occurring near SO trough (pink) and SO peak (sky blue) phases in the dominant hemisphere. **e**. Time course of V1-HC bias coherence relative to ripple onset (100ms sliding window with 20ms steps). **f**. V1-HC bias coherence for V1 bias from pre-onset window (-200 to 0s, left) and post-onset window (0 to 200ms, right). **g**. Distribution of V1 SO phases in dominant and non-dominant hemisphere at ripple peak. n = 42466 events, ρ = 0.34, p < 10^−300^, circular correlation. **h-i**. V1-HC reactivation bias coherence for events with SO trough phases in the dominant hemisphere only (light pink) and SO trough phases in both hemispheres (dark pink). **h**. Time course of V1-HC bias coherence relative to ripple onset (100ms sliding window with 20ms steps). **i**. V1-HC bias coherence for V1 bias from pre-onset window (-200 to 0ms, left) and post-onset window (0 to 200ms, right). For **e** and **h**, the shaded regions indicate the 95% bootstrapped confidence interval (CI) of the selected ripple events across all sessions. The shaded region in gray indicates 95% CI distribution of the event label shuffle. Colored bars above indicate time windows of significance where the 95% CI of the observed data exceeded the 95% CI of the shuffled distribution. For **f** and **i**, error bars represent the 95% bootstrapped CI of the selected events or the 95% CI of the event label shuffled data. Asterisks (*) denote significant coherence where the 95% CI of the real data does not overlap with the 95% CI shuffled distribution.

We found that ripple events occurring near the SO trough phase in the dominant hemisphere showed higher coherence between V1 and hippocampus. For ripple events occurring near the SO trough phase, V1-HC coherence was significantly elevated above the shuffled baseline across extended windows, both preceding and following ripple onset (−140 to 200 ms, 280 to 360 ms, 400 to 540ms; Fig. 4e). In contrast, those occurring near the SO peak phase showed significant reactivation coherence only within a brief window following ripple onset (60 to 120 ms; Fig. 4e). This phase-dependent difference was further confirmed by examining the coherence within the pre-onset (−200 to 0 ms) and post-onset (0 to 200 ms) windows separately: trough-phase events showed coherence significantly above the shuffled distribution in both windows, whereas peak-phase events either overlapped with the shuffled distribution (pre-onset) or were significantly reduced relative to trough-phase events (post-onset) (Fig. 4f). The results suggested that the reactivation coherence between V1 and hippocampus was higher when ripples occurred near the SO trough phase where V1 neurons in the dominant hemisphere were in a more active state.

While slow-oscillations in the dominant hemisphere influence V1-HC reactivation coherence, does the state of non-dominant hemisphere matter? We hypothesized that if there was competition between cortical ensembles representing track L and track R, asymmetries in slow-oscillations can further enhance the coherence of reactivating the memory in the dominant hemisphere. While SO activity across both hemispheres is often synchronous^30^, SO asymmetry can occur depending on sleep depth and neuromodulatory tone^31^. Similar to these observations^30,31^, our data showed a high correlation between left and right V1 SO phases at the ripple peak (ρ = 0.34, p < 10^−300^, circular correlation, n = 42466; Fig. 4g). However, we also observed many ripples with asymmetry in SO phases (n = 14670 out of 42466 ripples, ∼35%; Fig. 4g; i.e. one hemisphere was relatively closer to the trough phase while other hemisphere was relatively closer to the peak phase). When the SO trough was unique to the dominant hemisphere, V1-HC coherence was significantly elevated above the shuffled baseline across wide windows both before and after ripple onset (−160 to 200 ms, 340 to 360 ms, 400 to 420 ms; Fig. 4h). When both hemispheres were simultaneously near the SO trough, significant reactivation coherence emerged only following ripple onset (60 to 180 ms; Fig. 4h). This effect of hemispheric asymmetry was further confirmed by separately examining the pre-onset (−200 to 0 ms) and post-onset (0 to 200 ms) windows: V1-HC coherence exceeded the shuffled distribution in the pre-onset window only when the SO trough was unique to the dominant hemisphere, whereas post-onset coherence was significantly elevated for both conditions (Fig. 4i). Collectively, these results suggest an optimal SO phase relationship between the dominant and non-dominant hemispheres (being in trough and peak phase, respectively), during which the content of the hippocampal reactivation was most likely to match the content of pre-ripple cortical activity (Fig. 4h,i).

### Distinct and non-redundant contribution of ripple power, spindle-band power and slow-oscillations phase on cortico-hippocampal reactivation coherence

Having separately characterized how ripple power, spindle-band power, and SO phase each modulate V1-HC reactivation coherence, one outstanding question is whether these oscillatory features can jointly contribute to reactivation coherence or share a common underlying process. To address this, we constructed a unified Generalized Additive Mixed Model (GAMM) that simultaneously modelled all three features as predictors of reactivation coherence, allowing us to assess their independent predictive power and characterize their relationships with reactivation coherence. To estimate per-event reactivation coherence for GAMM, we calculated the signed geometric mean of HC and V1 track bias within a peri-ripple window (-100 to 100ms).

All three oscillatory features significantly predicted re-activation coherence within the same model (Fig. 5; Extended Data Fig. 7), indicating that their contributions are distinct and non-redundant. Ripple power (RMS of partial effect = 0.00903, p = 0.00322, n = 40259) and V1 spindle-band power in the dominant V1 hemisphere (RMS of partial effect = 0.00782, p = 0.0372, n = 40259) each acted as approximately linear predictors, with their partial effect on coherence increasing monotonically with power (Fig. 5a,b). In contrast, non-dominant V1 spindle-band power showed no significant relationship with coherence (RMS of partial effect = 0.00517, p = 0.500, n = 40259; Extended data Fig. 7a). For slow-oscillations, we observed bilaterally interacting oscillatory dynamics governing V1-HC coherence (Fig. 5c). Both the main effects of dominant and non-dominant hemisphere phase and their interaction were significant predictors of coherence (dominant: RMS of partial effect = 0.0130, p = 2.61×10^−6^, n = 40259; non-dominant: RMS of partial effect = 0.0132, p = 4.48×10^−6^, n = 40259; interaction: RMS of partial effect = 0.0158, p = 4.06×10^−4^, n = 40259;Extended data Fig. 7b). The total effect of SO dominant and non-dominant phase revealed that coherence was highest when the dominant hemisphere was near the SO trough while the non-dominant hemisphere was transitioning toward or already near the SO peak (Fig. 5c). More broadly, coherence was elevated when the dominant hemisphere was at an earlier phase relative to the non-dominant hemisphere during the trough phase window (π/2 to 3π/2), suggesting that SO phase asymmetry resulted in an optimal condition for coherent memory reactivation. These results remained similar when extending the analysis window from −100 to 100 ms to −200 to 200 ms (Extended data Fig. 8). Together, these findings demonstrate that ripple power, spindle-band power, and SO phase each make a distinct, non-redundant contribution to cortico-hippocampal reactivation coherence in which coherence is maximally enhanced when ripple power is high, the dominant hemi-sphere exhibits high spindle-band power and is in the trough phase of SOs, and the non-dominant hemisphere is in the peak phase.

**Figure 5.**
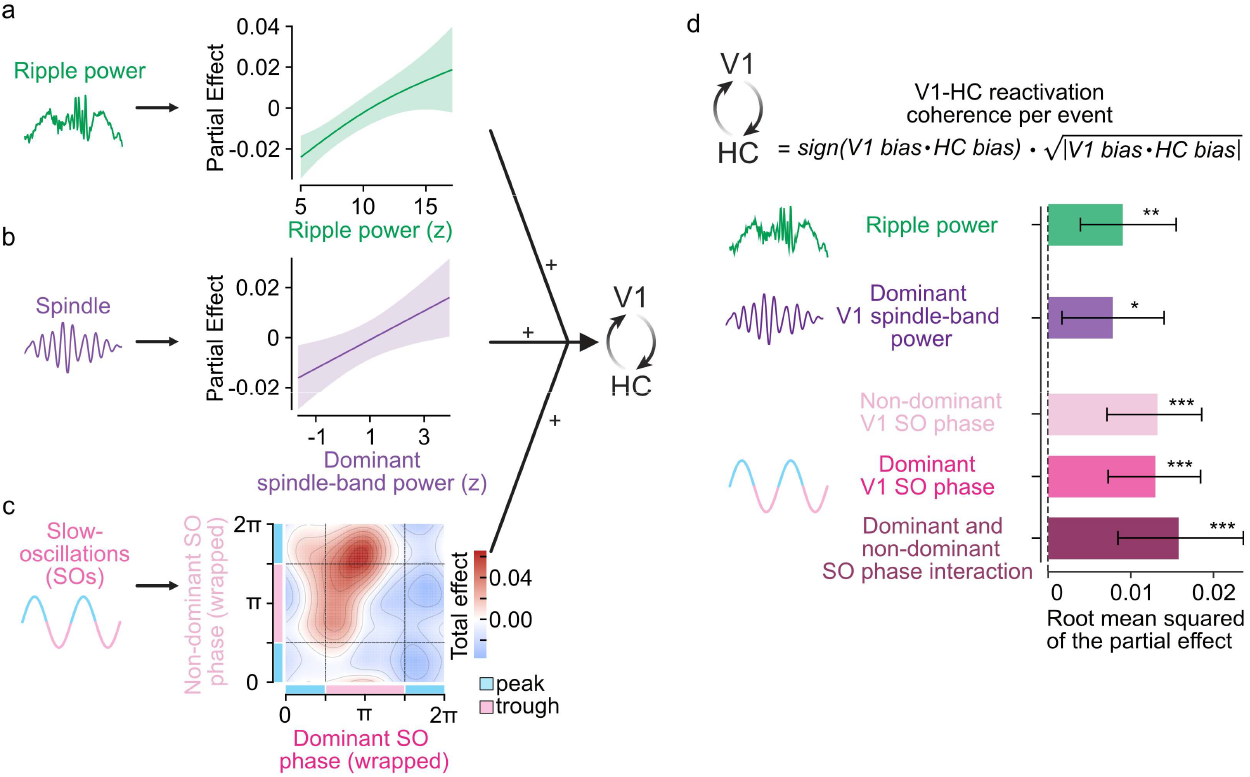
Distinct and non-redundant contribution of ripple power, spindle-band power, and slow-oscillation phase to V1-HC reactivation coherence. A Generalized Additive Mixed Model (GAMM) was constructed with hippocampal ripple power, V1 spindle-band power from dominant hemisphere, and V1 slow-oscillations (SOs) phase from both hemispheres (individual term as well as interaction term) simultaneously included as predictors of per-event V1-HC reactivation coherence. Reactivation coherence per event was estimated as geometric mean of V1 bias and HC bias (*sign*(*V1 bias* ⋅ *HC bias*) ⋅ 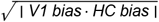). **a**. Partial effect of hippocampal ripple power on V1-HC reactivation coherence. Shaded region indicates the 95% confidence interval. **b**. Partial effect of V1 spindle-band power in the dominant hemisphere on reactivation coherence. Shaded region indicates the 95% confidence interval. **c**. Total effect of V1 slow-oscillation (SO) phase in the dominant (x-axis) and non-dominant (y-axis) hemisphere on reactivation coherence, shown as a bivariate heatmap. Phase windows corresponding to the SO peak and SO trough are indicated by green and pink shading on the respective axes. **d**. Root mean square (RMS) of partial effect for each predictor of the GAMM. Bars show the median bootstrapped RMS; error bars show the 95% bootstrapped confidence interval. Asterisks denote significance of each predictor in the model (\**p* < 0.05, \*\**p* < 0.01, \*\*\**p* < 0.001).

## Discussion

The brain faces a significant challenge during memory consolidation: from the many behavioural episodes encoded throughout the day, specific memories must be selected and reactivated simultaneously across distributed hippocampal and cortical circuits, spanning multiple brain regions. Our results advance understanding of this coordination on two fronts. First, content-selective cortico-hippocampal reactivation is maximally facilitated when three oscillations act in concert: 1) high ripple power in the hippocampus, 2) high spindle power localized to the reactivating memory in cortex (i.e. dominant hemisphere) and 3) a slow oscillation phase, particularly when the dominant neural circuit in the reactivation is in the SO trough phase, while the non-dominant neural circuit is in the SO peak phase. Second, this re-activation is not passively triggered by hippocampal ripples alone. Pre-ripple cortical activity already carries context-specific information that biases the subsequent hippocampal reactivation, suggesting that the cortex plays an active role in driving the content of memory selected for consolidation.

Traditionally, the hippocampus has been viewed as the primary driver of memory consolidation during non-REM sleep, using sharp-wave ripples together with the reactivation of recent memory traces as the teaching signal to drive content-specific reactivation in cortex, leading to their consolidation and long-term storage^1–3^. Slow oscillations and spindles are prominent during non-REM sleep and can occur co-nested with ripples, but have largely been viewed as providing desirable but content-agnostic windows to gate cortical responsiveness to hippocampal signal^1,3,8,10^. Our results challenge this division of labor. We show that the co-ordination of hippocampal ripples and local cortical oscillations together provides a spatio-temporally localized window for content-selective cortico-hippocampal communication. Reactivation coherence between cortex and hippocampus is maximally enhanced when all three act synergistically. By restricting optimal condition for cortico-hippocampal communication to selected cortical ensembles at a time^32,33^, this coordination may help to gate cortical dendritic plasticity^6,8,9,15^ in a content-specific manner, preventing spurious associations between concurrently active but competing memory traces from being consolidated together. This content-selective cortical plasticity may be instrumental for developing and stabilizing spatial modulation of visual representations observed in the visual cortex ^34–36^. This spatial modulation could arise from modifying local circuits, or increasing the influence of top-down inputs to visual cortex^37^, both of which could lead to the increased coherence of representations observed in visual cortex and hippocampus during behaviour^35,38^

Another important question is how does the brain decide which memory to selectively reactivate, from multiple competing experiences? Our result suggests that pre-ripple cortical activity biases subsequent content of hippocampal reactivation during ripple. Prior work has observed that an increase in cortical activity typically precedes hippocampal reactivations^6–8,39^, and this activity can be predictive of the hippocampal memory trace that reactivates^7,39^. However, it is important to distinguish between a general modulation of cortical activity that can co-occur with sleep rhythms and content-specific cortical reactivation. Here, we provide the evidence that content-specific cortical activity before ripple onset predicts the content of the subsequent hippocampal ripple – an effect particularly pronounced during asymmetries in local slow oscillations. This supports the view that competition between cortical ensembles before ripples can bias the reactivation content of the subsequent hippocampal ripple for memory prioritization^7,19,40,41^.

What may be the functional implication of slow-oscillations and spindles in selective memory consolidation? Slow-oscillations that are known to travel across the cortex^30,31,42^, sometimes as traveling waves^42^, could provide a roving window of opportunity to engage in the consolidation process while spindles are typically localized^15,43^ and could act as a spotlight on a specific cortical region and the memories represented within. These cortical dynamics may account for the phenomenon of targeted memory reactivation (TMR), where presenting sensory cues during sleep can aid consolidation of specific memories^44,45^. Our results on cortical SOs are consistent with the findings in human subjects where TMR is more effective during cortical SO up phases but not SO down phases^44,45^. We propose that local SO up phases, particularly when coinciding with high local spindle power, define the optimal window for engaging content-specific cortical circuits in coordinated reactivation with the hippocampus. This predicts that TMR efficacy could be substantially improved by targeting cues not just to global SO phase, but to the spatiotemporally localized windows of high spindle power and SO trough in the cortical circuit encoding the cued memory. Beyond TMR cueing, the likelihood that a given memory trace is reactivated may be influenced by the nature of experience before sleep including its novelty^46,47^, recency^48^ or salience^47,49^. Such experience-dependent biases could interact with travelling slow oscillations and localized spindles to preferentially consolidate specific memories during sleep.

Together, these findings recast the role of sleep oscillations in memory consolidation. Rather than merely providing a permissive state for a hippocampal teaching signal, the coordinated interplay of ripples, spindles, and slow oscillations establishes a precisely targeted, content-specific window for bidirectional cortico-hippocampal communication, where cortical dynamics actively shape which memories re-activate and consolidate. This shifts the focus of memory consolidation from the hippocampus alone to a dynamic dialogue between cortex and hippocampus, orchestrated by the oscillatory landscape of sleep.

## Acknowledgements

We thank members of the Institute of Behavioural Neuroscience (IBN) for valuable discussion. Mouse schematics in Fig. 1 and 2 were adapted and modified from SciDraw.io (http://doi.org/10.5281/zenodo.3925949, http://doi.org/10.5281/zenodo.3926215). This work was supported by the Medical Research Council scholarship (MR/N013867/1) to M.T.; Biotechnology and Biological Sciences Research Council Research grants (BB/T005475/1, BB/Z51665X/1, and BB/Y010345/1) to D.B.; and the Sir Henry Dale Fellowship from the Wellcome Trust and Royal Society (200501), the Human Frontier in Science Program (RGY0076/2018), Biotechnology and Biological Sciences Research Council grant (BB/W01579X/1), and UKRI Frontier Research grant (EP/Y024656/1) to A.B.S.

## Author contributions

This work was conceptualized by M.T., D.B. and A.B.S.; Experimental setup was by M.T., D.T. and E.A.B.H. Data collection was by M.T. Data processing was by M.T., D.T. and E.A.B.H; Methodology, software and formal analysis were by M.T.; Initial draft was written by M.T., D.B. and A.B.S.; Manuscript was revised and edited by all authors; Supervision and funding acquisition were by D.B and A.B.S. Both D.B. and A.B.S contributed equally. All authors have given approval to the final version of the manuscript.

## Methods

### Animal and surgery

For simultaneous chronic recordings in V1 and CA1, we used C57BL/6J wild-type mice (n = 6; four female and two male, age 16-27 weeks during recordings) obtained at around 4-6 weeks of age from Charles River UK Ltd. Mice were individually housed under a reversed 12-h light/dark cycle and experiments were performed during the dark phase of the cycle.

All animals were implanted with a custom-made circular headplate at ≥ 10 weeks of age under isoflurane anesthesia. Following 3-5 days of recovery, all animals were water restricted and trained to perform the virtual reality (VR) task and acclimate to a sleep box (30–60 min per day, 5–7 days/week). After about 5-8 weeks of VR training and sleep box habituation, animals underwent a second surgery under isoflurane anesthesia to perform craniotomies and chronically implant two Neuropixel probes, using a modified version of the Apollo drive^22^. We modified the inter-probe spacing to be 4.6mm (2.3mm from midline each). Two 1 mm craniotomies were made (AP:–3.4mm, ML: +/-2.3mm). Dura-gel (Cambridge NeuroTech), a biocompatible silicone, was applied to the exposed dura mater. We lowered the drive slowly (10µm/s) at an angle of 30° relative to the AP axis (tilted posterior-to-anterior), until a depth of around 3500µm, thus spanning all layers of V1, CA1 and dentate gyrus.

### Virtual reality environment and task

The two-track task involved mice running on two visually distinct familiar virtual linear tracks and licking left or right in the reward zone (95cm to 105cm) to receive a liquid reward. Two virtual linear tracks were designed and controlled using BonVision^50^, an open-source package for visual environment generation within Bonsai^51^. Each track was 140 cm in length and 8 cm in width and height. The ceiling, floor, and one-sided wall of the track were covered with a smoothed white-noise background and five visual landmarks that occupied only half of the display. The visual landmarks were 8 cm wide and centred at 30, 50, 70, 90, and 110 cm from the start of the track. Each track started with a single context-specific visual landmark at 30 cm, followed by four visual landmarks forming two visually repeating segments (segments from 34–61 cm and from 74–101 cm were visually identical; Fig 1a).

### Chronic electrophysiological recording

Electrophysiological recordings were performed chronically using two Neuropixels 2.0 probes^21,22^ (imec) from bilateral V1 and hippocampi. The neural signals were acquired via a PXI system (National Instruments) using SpikeGLX software (https://billkarsh.github.io/SpikeGLX/).

During the first 1–3 recording days, only visual stimuli (sparse noise and checkerboard patterns) were presented to facilitate identification of V1 and hippocampal layers and to optimize channel selection. Briefly, the earliest sink, based on current source density (CSD) analysis around the full-screen checkerboard-patterned stimulus onset, was used to identify V1 layer 4 (L4). The power spectral density (PSD) for each channel was calculated using a Hanning window of 4096 samples without overlap. V1 layer 5 (L5) can be identified based on the increase in the high-frequency power (300-600Hz) below L4. The CA1 region can be identified based on the increase in the ripple band power (125-300Hz) below L5 and the decrease in the theta power compared to the dentate gyrus below. All Neuropixels data were preprocessed with SpikeInterface^52^ (https://github.com/spikeinterface) with recordings from behaviour and sleep sessions from the same day concatenated prior to processing.. Spike sorting was performed using Kilosort 4^53^ (https://github.com/Mouse-Land/Kilosort).

Putative multi-unit activity (MUA) was obtained by pooling all non-noise clusters within each area. Putative single units were identified using quality metrics based on Bombcell^54^ (https://github.com/Julie-Fabre/bombcell) and SpikeInterface^52^. Clusters were classified as single units if they met:

1. *Waveform*: Mean waveform had a single trough and up to two peaks, with trough preceding and larger than peak.
2. *Spike amplitude*: Amplitude decreased with slope <–20µV/channel across sites.
3. *Trough-to-peak duration*: 0.1–0.8 ms.
4. *Waveform width*: Half-width of negative trough ≤0.3 ms.
5. *Amplitude cutoff*: ≤10% contamination from overlapping spikes.
6. *Sliding refractory period violations*: <10% (90% confidence, 0.25 ms bins).
7. *Amplitude variability*: CV and range ≤0.7.
8. *Mean firing rate*: ≥0.1Hz.
9. *Cluster metrics*: Standard deviation ratio ≤4; SNR ≥2. Importantly, only single units with stable spatial tuning for one or both tracks were included for further analysis.
10. *Spatial tuning*: The peak spatial firing of each cluster was greater than 95% of the shuffled peak distribution (i.e. Circular spike train shift within each lap).
11. *Spatial tuning stability*: The correlation between the odd-trial ratemap and the even-trial ratemap was greater than that of the 95% of the shuffled correlation distribution (i.e. Circular spike train shift within each lap).

### Sleep classification

Sleep stage was classified based on animal immobility and local field potentials (LFPs) recorded from V1 and hippocampus, modified from previous methods^7,55^. Animal movement was extracted from behavioral camera recordings (30Hz, Raspberry Pi HQ camera) and quantified as the magnitude of summed pixel changes between consecutive frames. The immobility threshold was determined by visual inspection of the movement distribution. Putative sleep epochs were defined as immobility periods lasting longer than 10s^7^. Short interruptions in immobility were tolerated, such that movement bouts shorter than 1.5s did not terminate a sleep epoch. Non-rapid eye movement (NREM) sleep was detected using k-means clustering of the V1 SO-band (0.5–4Hz) power, yielding epochs dominated by high SO activity. For sleep epochs not classified as NREM, rapid eye movement (REM) sleep was identified based on the ratio of hippocampal theta (6–9Hz) to delta (0.5–4Hz) power. Epochs were classified as REM when the theta/delta ratio exceeded 1 and followed a preceding NREM period. Transitions from NREM to REM were constrained such that the maximum allowable interval between NREM to REM was 120 s. Only REM sleep epochs longer than 10 s were included for further analysis. In addition, immobility epochs longer than 2 s that did not meet criteria for NREM or REM were classified as quiet wake. Within these quiet wake periods, small movements shorter than 0.5 s were tolerated. This state corresponds to immobility without the electrophysiological signatures of sleep. All subsequent ripple detection and reactivation analysis were restricted to NREM window only.

### Local field potential processing and Sharp-wave ripple detection

LFP from a putative CA1 channel with highest ripple power or a putative layer 5 V1 channel from each hemisphere was band-pass filtered based on the frequency range of interest (0.5 – 4Hz for slow oscillations, 9-17Hz for spindle-band oscillations and 125-300Hz for ripple-band oscillations).The band-pass filtering was performed using with a finite impulse response (order = (6 × sampling rate)/(filter frequency width)) zero phase filter. The z-scored instantaneous power or phase of the filtered LFP signal was calculated using a Hilbert transform.

Ripple detection was based on a modified version of a previously used algorithm^56^ (Extended data Fig. 4). Event onset and offset thresholds were set to a z-scored ripple power of 2. Events with an inter-ripple interval shorter than 30 ms were merged. A putative ripple event was accepted if its peak ripple power exceeded 5 and its duration was between 30–200 ms. Due to the bilateral synchrony and the presence of ripple chains, we removed events with inter-ripple intervals less than 50ms, which resulted in 42,466 ripple events (mean±SE =1930 ± 136.1).

### Track decoding based on Partial least squares regression and kernel density estimation

To quantify track decoding in V1 and hippocampus during behavior and during reactivation, we optimized this decoding process using three step approach for improved track discrimination: 1) We used partial least squares regression^57,58^ to extract optimal latent variables that maximized the covariance between the track identity and population spiking activity, such that the representation of the two tracks could be better separated, 2)we then used kernel density estimation (KDE) to estimate the probability density function of each track’s representation by the latent variables. 3)Finally, to measure which track was currently represented in the neural population, we calculated a log-odds metric^5–7^. This metric represented track identity and likelihood by its sign and magnitude, respectively (e.g. large positive value-high likelihood of Track L, small negative value-low likelihood of Track R).

#### Partial least squares regression

We first constructed the matrix of spike counts (100ms, z-scored relative to the entire session) from all spatially tuned cells across both hemispheres. We then extracted the three PLS latent variables that maximized the covariance between binned z-scored population spike counts and track identity. Mathematically, the objective of PLS is to find weight vectors **W** for the neural data and weight vectors **Q** for the track identity label that maximize the covariance between the projected neural activity and track label:

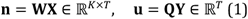

where ***K*** denotes number of latent variables (K = 3), ***T*** denotes number of time bins, **X** ∈ ℝ^*N×T*^represents the spike counts matrix, ***N*** denotes number of neurons, **n** represents the projection of the neural activity **X** onto **W, Y** ∈ ℝ^*T*^ represents the track identity matrix, and, **u** represents the projection of the track identity **Y** onto **Q**.

For the k^th^ component, this optimization can be written as:

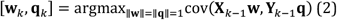

where **X**_*k-1*_ and **Y**_*k-1*_ are the residual matrices after deflating the first *k* −1 components.

The neural weight vectors **w**_***k***_ were then converted into orthonormalized loading vectors **P**_***k***_, which quantify how strongly each neuron contributes to a latent component *k*:

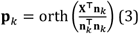

where **P**_***k***_ denotes the orthonormalized loadings (each column unit length and mutually orthogonal).

Thus, the orthonormalized loadings **P** form the projection matrices, allowing the transformation of high-dimensional spike-count vectors into a low-dimensional PLS latent space:

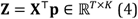

where **Z** is the PLS latent representation of population activity, with each row corresponding to the activity pattern at each time bin.

#### Kernel density estimation

Next, we applied kernel density estimation (KDE) on PLS-projected vectors **Z**, allowing us to calculate the relative likelihood of a Track L versus Track R representation given a pattern of neural activity. For a given track 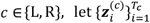 denotes the set of PLS-projected vectors when animal was running on a given track *c*, where *T*_c_ represents the total number of time bins for that track.

The probability density of observing each PLS-projected vector *z* under the distribution of track ***c*** representation was estimated as:

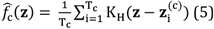

where *K*_H_ is a multivariate Gaussian kernel with bandwidth matrix *H*. The optimal bandwidth was determined empirically by maximizing the separation between track distributions in the latent space using receiver operating characteristic curve with 10-fold cross-validation (see below).

#### Log odds metric

To quantify the relative likelihood of a Track L versus Track R representation (i.e. track bias) at each time bin, we calculated the log odds between Track L and Track R KDE likelihoods:

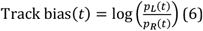

where *p*_L_(*t*) and *p*_R_(*t*) denote the KDE probabilities of Track L and Track R at time *t*, respectively.

The reactivation track bias, therefore, was quantified as a function of time from ripple onset for all ripple events (-1 to 1 second, 20ms bins with 10ms step). Due to the high synchrony between left and right hippocampal ripples, ripple events occurring within 50 ms of each other were treated as duplicates, and only the first event was retained for subsequent analysis. To further ensure the independence of the track bias measure between ripple events, any bins overlapping with the preceding or subsequent ripple events were excluded. The track bias at each time bin were z-scored relative to the track bias distribution over the entire sleep session. This track bias measurement allowed us to identify track-biased ripple events based on the mean HC track bias during each individual ripple, where the positive values corresponded to Track L bias and the negative values to Track R bias. A similar track bias metric has been used previously^20,59,60^ to quantify relative bias between two contexts during behavior or reactivation.

#### Inclusion of position bins with above-chance track discriminability during behaviour for reactivation decoding during sleep

For track decoding during VR task, we used all spiking data when animals were locomoting on Track L and Track R (> 1 cm/s). For reactivation track decoding during sleep, to ensure that our reactivation decoding reflected context-specific features rather than shared features, the PLS components used for ripple projection were constructed using spiking data from only position bins with above-chance track discriminability. We identified position bins with poor track discriminability during VR based on logistic ridge regression with 10-fold cross-validation.

For a given spatial bin i, the probability of the track being correct (y_i_ = 1) given the projected data z_i_ is modelled as:

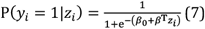

where the coefficients *β* are found by a cost function that minimizes the negative log-likelihood with an L2 penalty term:

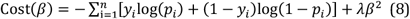

where λ is the regularization parameter selected based on 10-fold cross-validation.

The decoding accuracy at each position bin was then quantified as the proportion of correctly classified time bins. Control accuracies were obtained from 1000 iterations of Cell-ID shuffling. Model fitting and evaluation were performed based on the same 10-fold cross-validation framework: the ridge penalty λ was selected on the training folds, and decoding accuracy was computed on the held-out fold. The final bin-wise accuracies were averaged across all 10 folds. Position bins were excluded if decoding accuracy fell within the 95% of the shuffled distribution (Extended Data Fig 3).

### Receiver operating characteristic curve for track discriminability during behavior

Overall discriminability between the two tracks during VR task was quantified using the area under the receiver operating characteristic curve (AUC). The ROC curve was constructed by thresholding the track bias at different levels, and AUC was defined as:

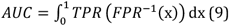

where *TPR* and *FPR* are the true and false positive rates at different track bias threshold, respectively. *AUC* was computed with 10-fold cross-validation where AUC = 0.5 indicates chance-level discriminability.

Statistical significance was assessed by comparing the distribution of AUC values from the real data (1000 bootstrap iterations) with a null distribution obtained from track label shuffling (1000 iterations). For all analyzed sessions, the 95% confidence intervals of the real-data AUC exceeded those of the shuffled data.

### Reactivation coherence analysis

To quantify reactivation coherence between V1 and HC during sleep, we first selected Track L and Track R biased ripple events based on mean track bias calculated from region A within a specified time window relative to ripple onset (e.g. 0 to 100ms in HC) followed by calculating the track bias difference between Track L-biased and Track R-biased ripple events in region B (e.g. -200 to 0ms in V1). To ensure the results were not driven by an arbitrary track bias threshold for event selection, we systematically shifted the track bias threshold (i.e. 10 deciles) for event selection and examined how the mean track bias difference at different times relative to ripple onset in region B varied with the track bias thresholds in region A. The V1-HC reactivation bias coherence for a given time window was then summarized as the area under the track bias difference curve (AUC).

To characterize the temporal dynamics of reactivation bias coherence relative to ripple onset, we computed bias coherence using a 100 ms sliding window with 20 ms steps. At each time bin, the observed bias coherence was considered significant if its 95% bootstrapped confidence interval did not overlap with the 95% confidence interval of a null distribution derived from 1,000 iterations of event label shuffling. The window of significant reactivation coherence was defined as a contiguous run of at least two consecutive significant time bins.

For determining the statistical significance of the overall reactivation coherence before and after ripple onset, we compared the bootstrapped distribution (1000 bootstrap iterations) of the bias coherence values calculated from the time windows before (-200 to 0ms, pre-onset) and after (0 to 200ms, post-onset) ripple onset with a null distribution obtained from event label shuffling (1000 iterations). The reactivation bias coherence was considered significantly above the baseline if the 95% boot-strapped confidence interval did not overlap with the 95% confidence interval of the shuffled distribution.

For the analyses in Fig. 2, we identified Track L and Track R biased events based on HC track bias during the ripple period (0 to 100ms) and/or ripple power. We then quantified the track bias difference in V1 at different times from ripple onset.

In contrast, for Fig. 3 and 4, events were selected based on both V1 track bias and specific V1 oscillatory features (e.g., spindle-band power or SO-band phase) from either the dominant or non-dominant V1 hemisphere — where the dominant V1 hemisphere is defined as the one associated with the reactivation track bias in V1. We then examined the resulting track bias difference in the HC during the ripple window (0 to 100ms). This reversal in selection criteria was necessary to investigate how V1 oscillatory features influence V1-HC reactivation coherence without introducing circular reasoning. Because V1 spiking activity is coupled to the V1 LFP activity, selecting events based on HC track bias and then partitioning them by V1 oscillatory features would have created an inherent bias. By selecting events based on both V1 track bias and V1 oscillatory features, we ensured a more rigorous quantification of V1-HC reactivation coherence.

### Linear mixed-effect model for track-selective ripple modulation of V1 neurons

To determine whether V1 neurons were modulated by ripples in a track-selective manner (as defined by HC reactivation track bias) across different time windows relative to ripple onset (e.g.-200 to 200ms, -200 to 0ms or 0 to 200ms), we used a linear mixed-effects regression model. For each V1 neuron, we first computed the normalized firing-rate difference (Δ*FR*_track_) between Track L (*FR*_Track L_) and Track R (*FR*_Track R_) during the VR task:

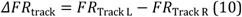

We then computed the normalized firing-rate difference (Δ*FR*_ripple_) be-tween Track L–biased ripple events (*FR*_Track L ripple_, top 25% positive HC track bias) and Track R–biased ripple events (*FR*_Track R ripple_, top 25% negative HC track bias):

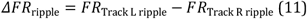

We then applied a linear mixed-effects regression model to quantify whether each V1 neuron’s track-selective firing-rate difference during behavior (Δ*FR*_track_, fixed effect) predicts the track-selective firing-rate difference during ripples (Δ*FR*_ripple_, response variable), while accounting for the random effect due to animal and session variability:

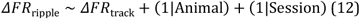

where (1|Animal) and (1|Session) terms are random intercepts grouped by animal ID and session ID.

Each predictor was considered significant if the 95% confidence interval of the *β* coefficient did not overlap with zero. A positive *β* coefficient would indicate that V1 neurons representing the same track reactivated in HC show corresponding increase in firing rate during ripples.

### Generalized additive mixed models for relationship between oscillatory features and V1-HC reactivation coherence

To examine how hippocampal ripple power, V1 spindle-band power and V1 SO phase at the time of ripple contributed to V1-HC reactivation coherence, we constructed Generalized additive mixed models (GAMM) that simultaneously modelled all three oscillatory features as predictors of re-activation coherence while accounting for the random effect due to animal and session variability.

To obtain a single per-event coherence estimate suitable for GAMM, we calculated the signed geometric mean of V1 and HC track bias within a peri-ripple window of −100ms to 100ms:

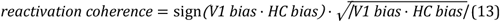

This coherence metric captures both the magnitude and the sign of co-reactivation, with positive values indicating coherent reactivation of the same track across V1 and HC.

All GAMMs were fit using the bam() function from the mgcv package in R, with fast restricted maximum likelihood for parameter estimation and covariate discretization enabled. The model (Fig. 5) was specified as:

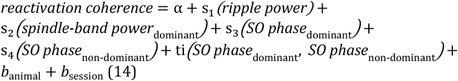

where s_1_ and s_2_ are penalised regression splines (basis dimension k = 5) capturing the smooth partial effects of z-scored ripple power and dominant-hemisphere V1 spindle-band power, respectively; s_3_ and s_4_ are cyclic cubic regression splines (k = 8) capturing the main effects of dominant- and non-dominant-hemisphere SO phase; and ti(⋅,⋅) is a tensor-product interaction spline (cyclic cubic basis, k = 8) capturing variance in coherence attributable to the interaction between dominant and non-dominant SO phase beyond their main effects. Cyclic splines were used for all SO phase terms to respect the circular nature of phase (−π to π). Animal and session were included as random intercepts 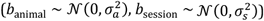 to account for between-subject and between-session variability. Significance of smooth terms was assessed via the *p*-values estimated from the penalized likelihood ratio test.

Furthermore, four complementary effect size metrics were estimated for each smooth term via a bootstrapping procedure (1000 iterations). For each iteration, data were resampled with replacement from the full dataset and the model was refit.

Partial deviance explained was computed using a drop-one approach, where a reduced model was refit with the target term omitted and the partial deviance taken as the difference in percentage deviance explained between the full and reduced models. Partial *η*^2^ (Eta squared) was calculated 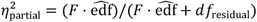, where *F*is the F-statistic, 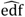, the degrees of freedom, and *df*_residual_the residual degrees of freedom.

Peak-to-trough amplitude and root mean square (RMS) were derived from the isolated partial effect of each smooth term, obtained by extracting the relevant columns from the linear predictor matrix and zeroing all others, with non-focal predictors held at their mean. Amplitude was defined as the range of this isolated prediction and RMS as the square root of its mean squared value. The bootstrapped median and 95% confidence interval were reported for each metric (Fig 5d; Extended Data Fig 7c, 8d).

To assess whether non-dominant hemisphere spindle-band power provided additional predictive value, we fit an extended model (Extended Data Fig 7) that added a further smooth term *s*_5_(*spindle-band power*_*non-dominant,ij*_)(*k* = 5) to the primary model (Fig 5; equation 13). As this term did not reach significance, it was excluded from the primary model. Finally, to verify robustness to the choice of coherence estimation window, we additionally repeated this extended model using per-event coherence calculated over a wider peri-ripple window of −200 to 200 ms (Extended Data Fig 8).

**Extended Data Fig 1.**
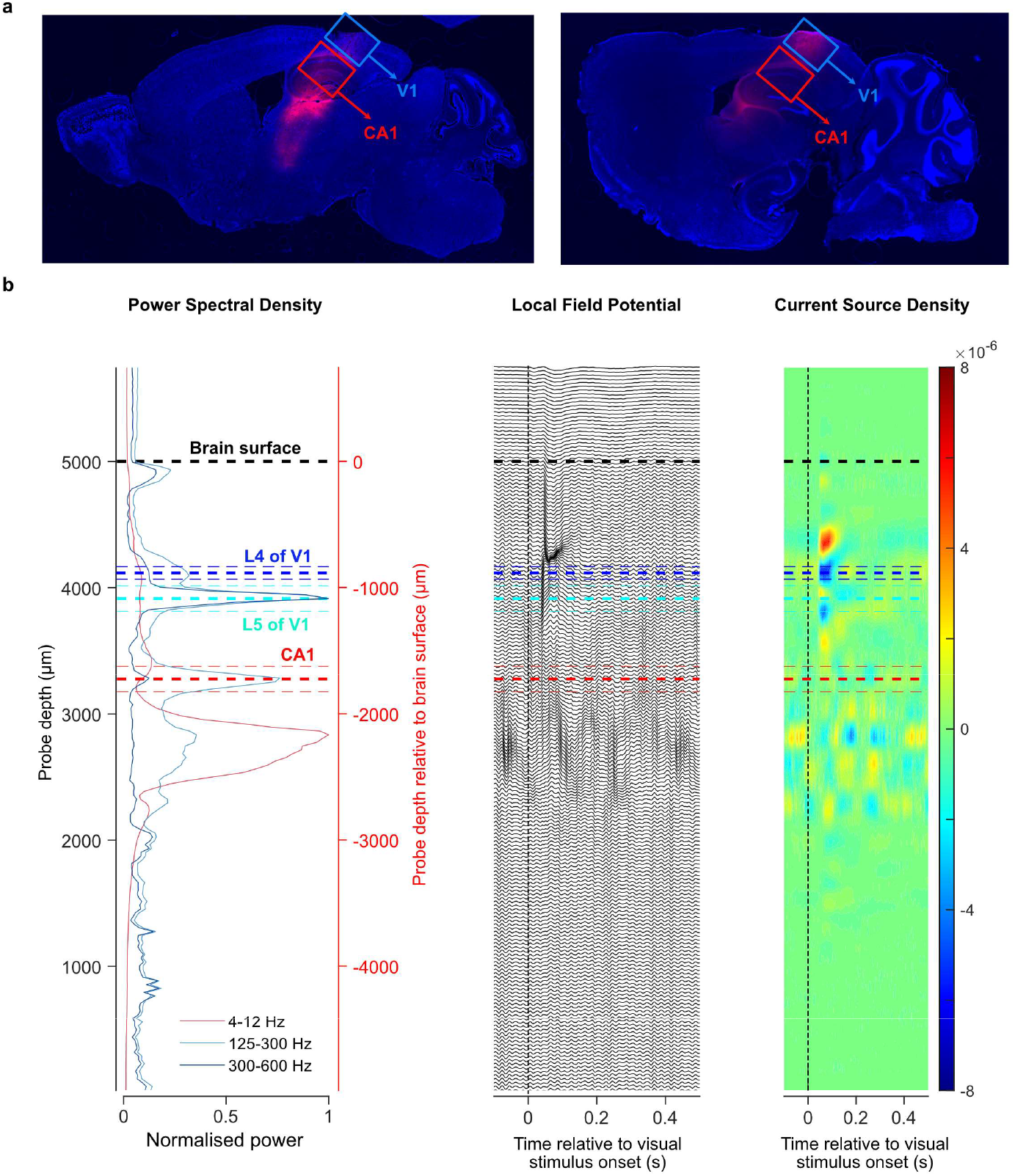
Histological and electrophysiological confirmation of V1 and CA1 region of HC. **a**. Example histology of probe trajectories through V1 and hippocampus for chronic recording (sagittal section). **b-d**. The local field potential (LFP) and current source density (CSD) depth profile from a Neuropixel 2.0 probe used to identify layer 4 (L4, dark blue) and layer 5 (L5, cyan) in V1 and the CA1 region of hippocampus (red). **b**. Normalised power spectral density depth profile of three different frequency bands: 4–12 Hz (theta, light red), 125–300 Hz (ripple, light blue), and 300 Hz and above (high-frequency band, dark blue). The Y-axis represents the probe electrode’s depth, where 0 indicates the tip of the probe. **c-d**. Averaged **(c)** LFP and **(d)** CSD depth profile during relative to visual stimulus onset (500ms checkerboard stimulus). L4 can be identified based on the earliest sink. L5 can be identified based on the peak in the high-frequency band below L4. The CA1 region can be identified based on the peak in the ripple band below L5. The Y-axis on the right indicates probe depth relative to the brain surface.

**Extended Data Fig 2.**
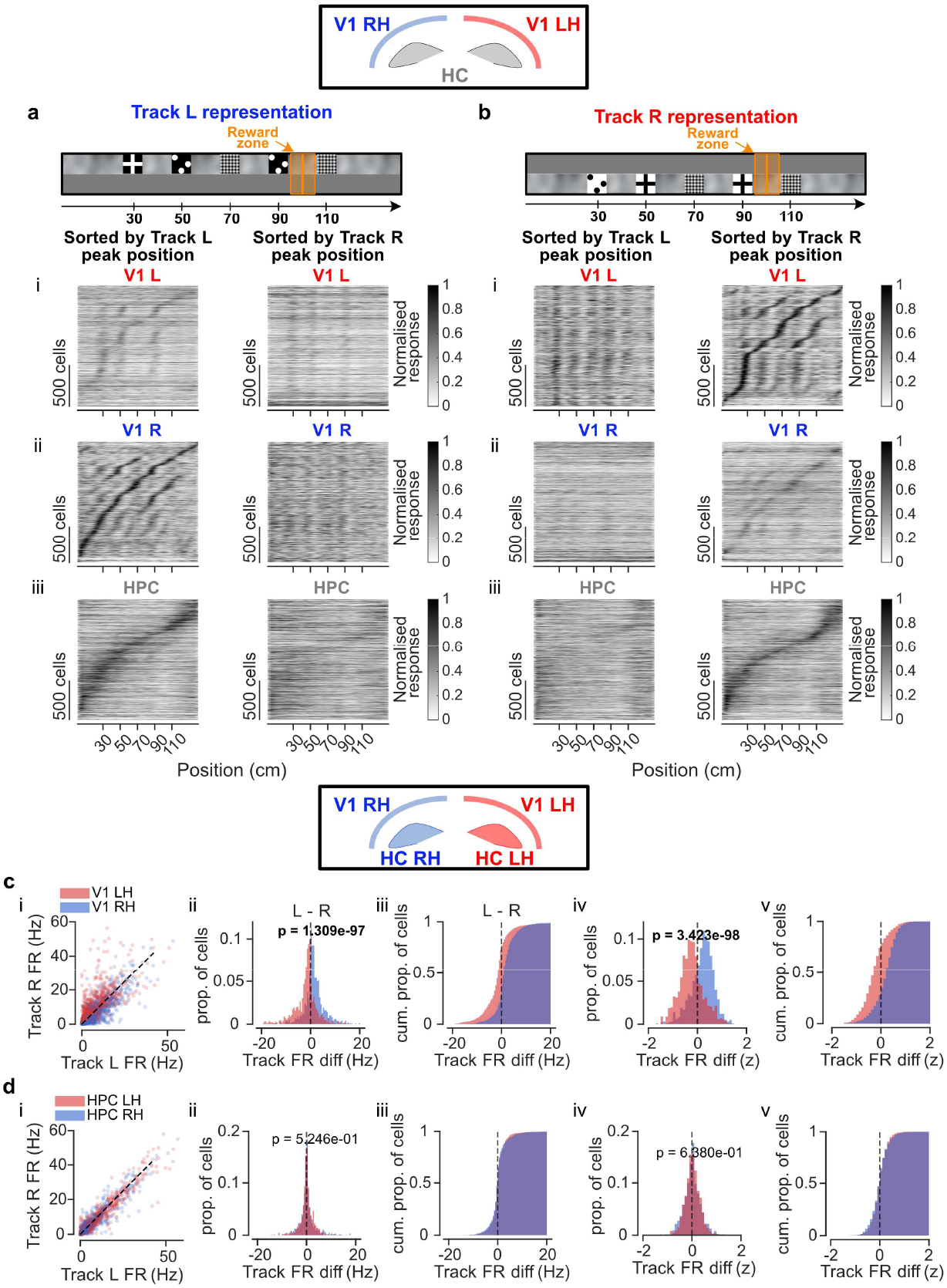
Context-selective spatial representations were hemispherically biased in V1 but not in the hippocampus. **a-b**, Normalised spatial response as a function of spatial position on **(a)** Track L and **(b)** Track R during even laps in (i) left V1, (ii) right V1, and (iii) both hippocampi, based on activities from odd laps. Cells were pooled across sessions and sorted according to the position of maximum activity during odd laps. **c-d**, Distribution of mean firing rate on Track L and Track R of neurons from left (blue) and right (red) **(c)** V1 and **(d)** hippocampus. (i) Mean firing rate on Track L vs Track R. (ii) Distribution of mean firing rate difference between Track L and Track R (Kolmogorov–Smirnov test: V1 LH vs V1 RH, p = 1.31 × 10^−97^; HC LH vs HC RH, p = 0.525). (iii) Cumulative distribution of mean firing rate difference between Track L and Track R. (iv) Distribution of mean z-scored firing rate difference between Track L and Track R (Kolmogorov– Smirnov test: V1 LH vs V1 RH, p = 3.42×10^−98^; HC LH vs HC RH, p = 0.638). (v) Cumulative distribution of mean z-scored firing rate difference between Track L and Track R.

**Extended Data Fig 3.**
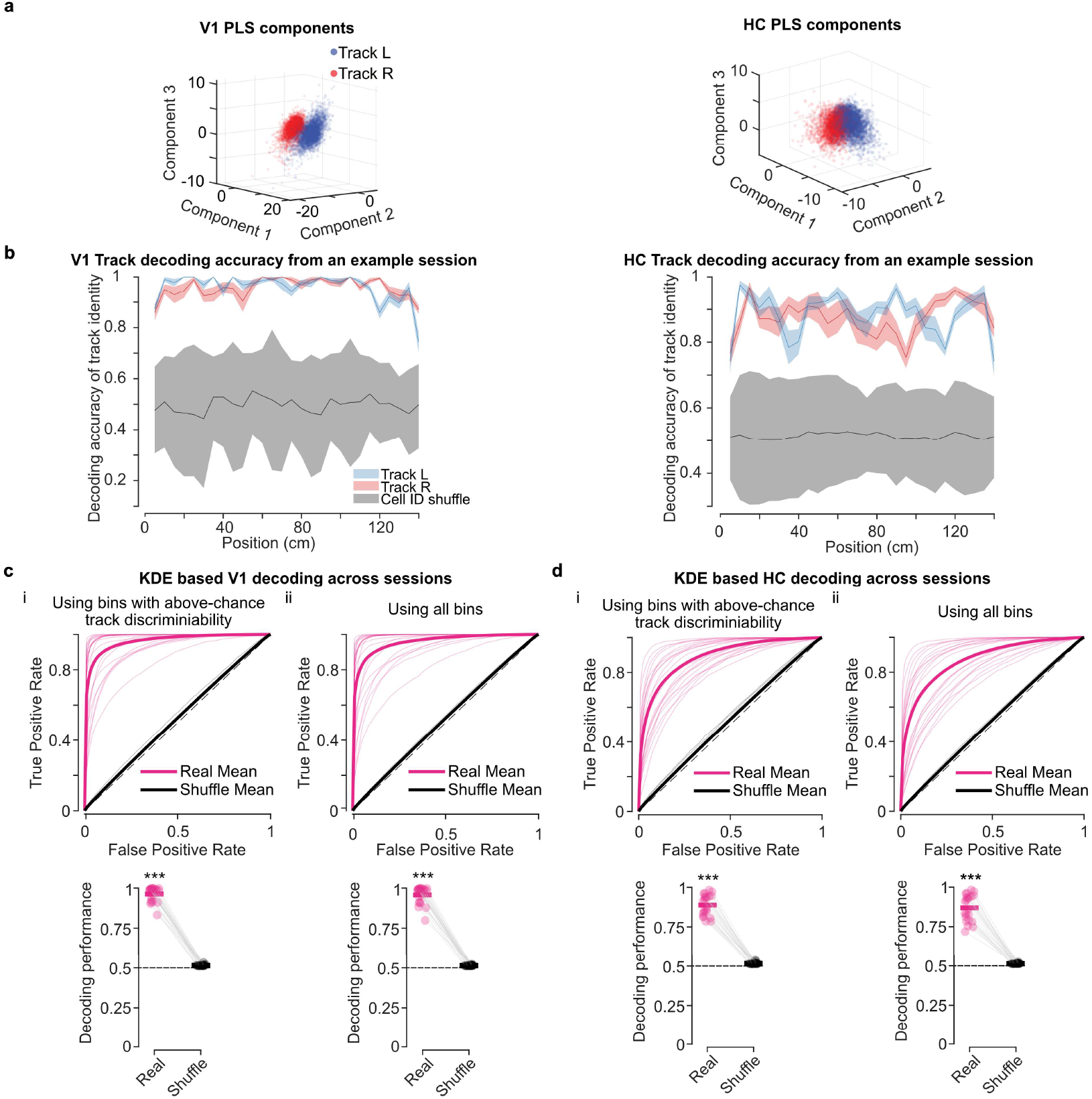
PLS based context decoding using all bins or only position bins with above-chance level track discriminability. Partial least square (PLS) regression was applied to obtain latent PLS components that best capture the covariance between population spike trains (100 ms bins) and track identity. **a**, Top three PLS latent components based on hippocampal (HC, left panel) and V1 (right panel) spike trains on Track L (blue) and Track R (red) from an example session. **b**, Decoding accuracy of track identity as a function of spatial position based on 10-fold cross-validated logistic regression using population activities from HC (left panel) and V1 (right panel) of an example session on Track L (blue) and Track R (red). Shaded regions of Track L and Track R decoding indicate standard error across 10 folds. Shaded regions of cell ID–shuffled data (grey) indicate the 95% confidence interval using 1000 bootstrap iterations. Position bins with decoding accuracy significantly higher than the shuffled distributions were considered positions bins with above-chance level track discriminability and included reactivation analysis. **c-d**, Track identity decoding based on kernel density estimation with 10-fold cross-validation where the decoding performance was quantified using the area under the receiver operating characteristic curve (ROC AUC). **c**, decoding using V1 spike counts (speed > 1cm/s) from (ci) position bins with above-chance track discriminability (AUC: real vs track-label shuffle 95th percentile, mean±SE = 0.963±0.00957 vs 0.516±9.7×10^−4^, Wilcoxon signed-rank test, ***p = 2.15×10^−5^) and (cii) all bins (AUC: real vs track-label shuffle 95th percentile, mean±SE = 0.957±0.0115 vs 0.515±5.64×10^−4^, ***p = 2.15×10^−5^). **d**, decoding using HC spike counts (speed > 1cm/s) from (di) position bins with above-chance track discriminability (AUC: real vs track-label shuffle 95th percentile, mean±SE = 0.888±0.0139 vs 0.517±0.00137, Wilcoxon signed-rank test, ***p = 2.15×10^−5^) and (dii) all bins (AUC: real vs track-label shuffle 95th percentile, mean±SE = 0.870±0.0178 vs 0.515±5.92×10^−4^, ***p = 2.15×10^−5^). For c-d, pink and grey are used to represent real and track-label shuffled data, respectively. Top panel represents ROC for each session and mean ROC across sessions. Bottom panel represents AUC for each session and mean AUC across sessions.

**Extended Data Fig 4.**
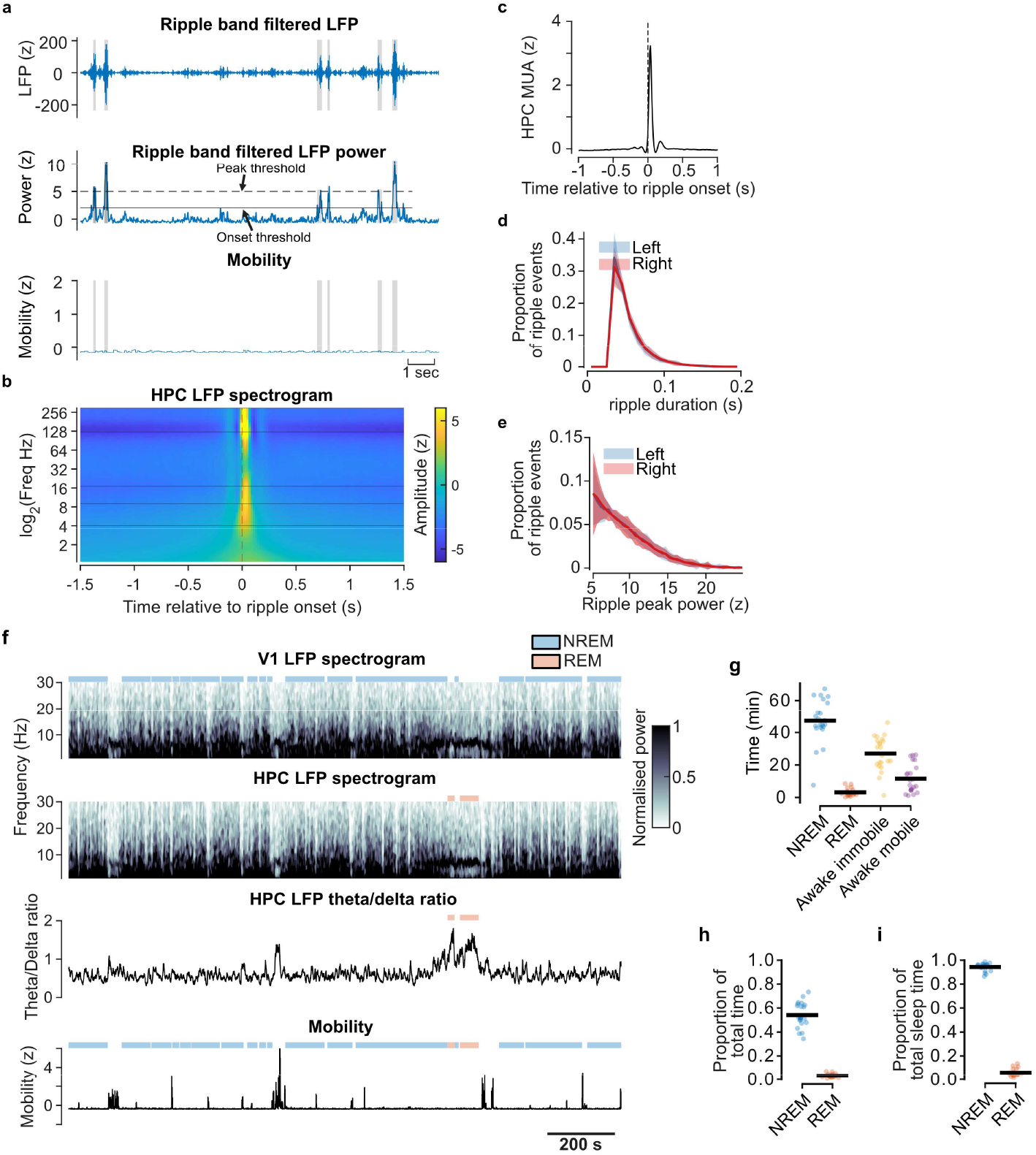
Detection of hippocampal ripple and sleep state. **a**. Example traces of ripple band filtered LFP (top), ripple band filtered LFP power (middle), and animal mobility (bottom, z-score of the pixel change) from an example session. The shaded regions in grey represent the detected ripples. **b**. Average hippocampal LFP spectrogram relative to ripple onset. **c**. Averaged z-scored hippocampal MUA activity relative to ripple peak. The shaded regions indicate standard error across sessions (n = 22 sessions). **d**. Distribution of ripple duration for left (blue) and right (red) hippocampus. The shaded regions indicate the standard error across all sessions (n = 22 sessions). **e**. Distribution of ripple peak power for left (blue) and right (red) hippocampus. The shaded regions indicate the standard error across all sessions (n = 22 sessions). **f**. Example session showing the V1 and hippocampal (HC) LFP spectrograms (0–30Hz), hippocampal theta/delta ratio, and mobility trace across time. Periods classified as NREM (blue) and REM (orange) are indicated above each trace. NREM epochs were identified based on high V1 delta (0.5–4Hz) power, while REM epochs were detected based on elevated hippocampal theta/delta ratio (ratio>1) following NREM. Mobility was quantified from behavioural video (30Hz) as the summed pixel change between consecutive frames, with immobility thresholds used to distinguish sleep from quiet or mobile wake. **g**. Average time spent in each state (NREM, REM, quiet wake, mobile wake) across sessions (mean ± SEM, dots = individual sessions). **h**. Proportion of total recording time spent in NREM and REM sleep. **i**. Proportion of total sleep time spent in NREM and REM. From **g-i**, individual data points represent single sessions; bars indicate the mean across all sessions (n = 22 sessions).

**Extended Data Fig 5.**
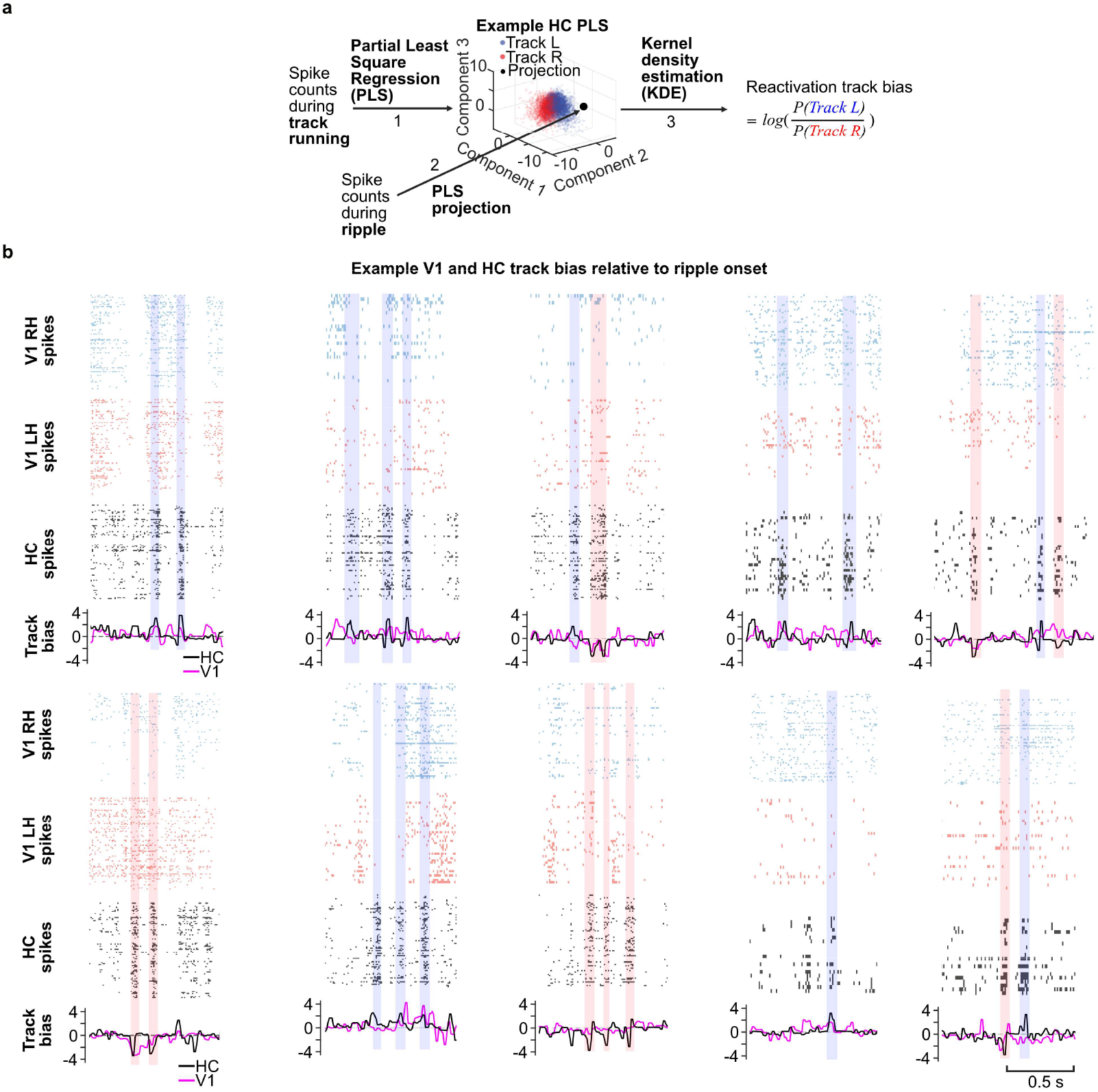
Decoded track bias in V1 and hippocampus during ripples. **a**. Simplified schematic diagram for the track decoding pipeline based on partial least square regression (PLS) and kernel density estimation (KDE): 1. Partial least square regression (PLS) was applied to extract top three latent components that maximised the covariance between z-scored V1 or HC population spike counts (100ms bin) and track identity when animal was running on Track L (blue) and Track R (red). The orthonormalized PLS loading matrices would then serve as projection matrices. 2. Following extraction of PLS components, z-scored spike counts (20ms bin with 10ms step) during ripples were projected onto PLS latent space. 3. Kernel density estimation (KDE) was applied to quantify the likelihood that a given projected neural pattern corresponded to a Track L or Track R representation. To measure relative likelihood of Track L versus Track R representation, we calculated log odds between Track L and Track R KDE likelihoods to infer reactivation track bias at each time bin. **b**. Example Spiking activity and track bias of V1 and HC from different reactivation events. Example Track L biased (shaded in blue) and Track R biased (shaded in red) ripple events are labelled based on HC track bias.

**Extended Data Fig 6.**
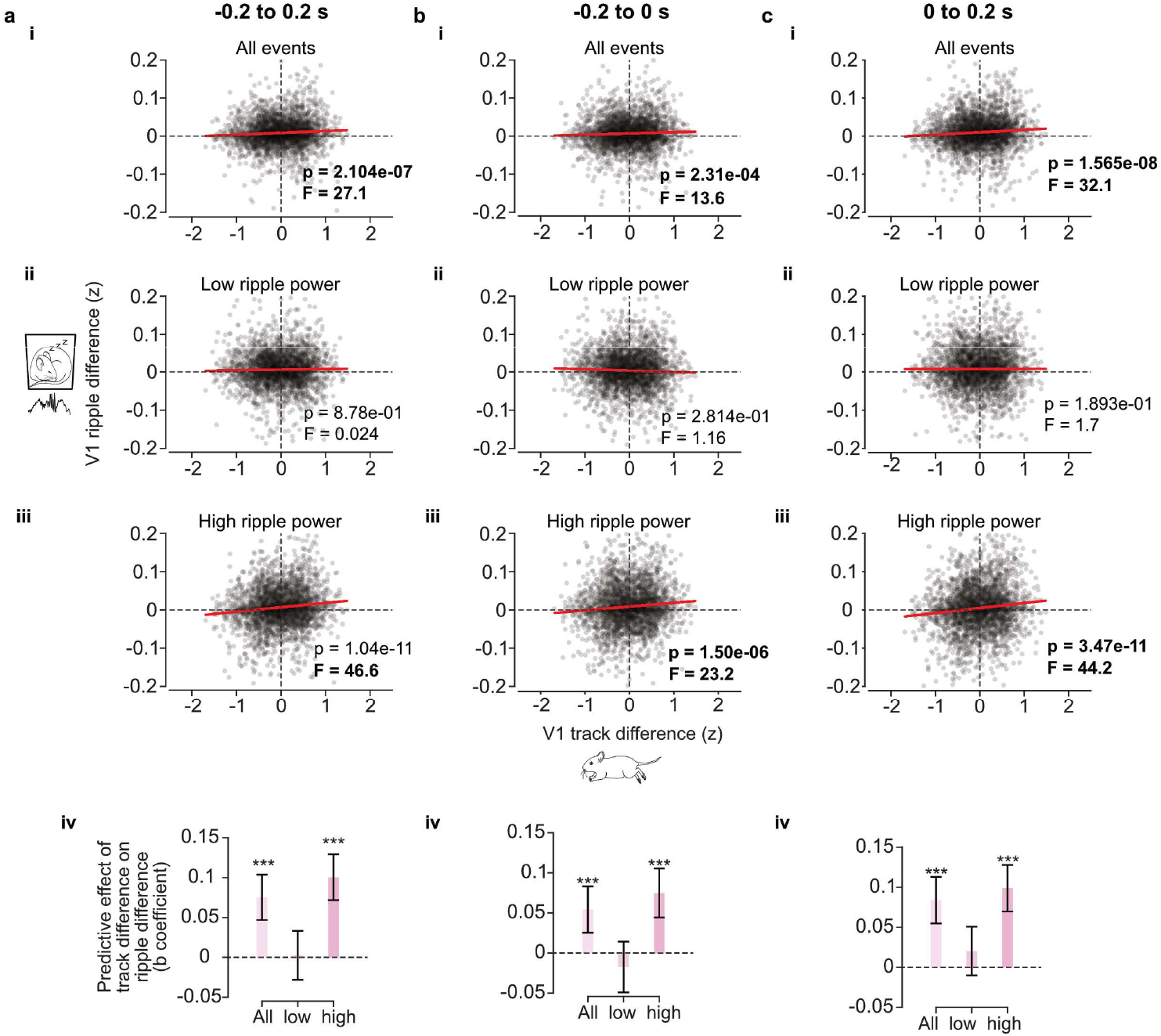
Context-selective ripple modulation of V1 neurons depends on ripple power. **a-c**. Linear mixed-effect regression for the relationship between V1 firing rate difference across Track L and Track R during VR and V1 firing rate difference across tracks during HC Track L and Track R reactivation within **(a)** −200 to 200 ms window before ripple, **(b)** −200 to 0 ms window before ripple and **(c)** 0 to 200 ms window before ripple. **(ai, bi, ci)** All events. **(aii, bii, cii)** Low ripple power events. **(aiii, biii, ciii)** High ripple power events. **(aiv, biv, civ)** Predictive effect (b coefficient) of each predictor (V1 activity during all ripples, low-power ripples and high-power ripples) using linear mixed-effect regression. Error bar indicate 95% confidence interval of the standardized b coefficient of the predictors.***p < 0.001.

**Extended Data Fig 7.**
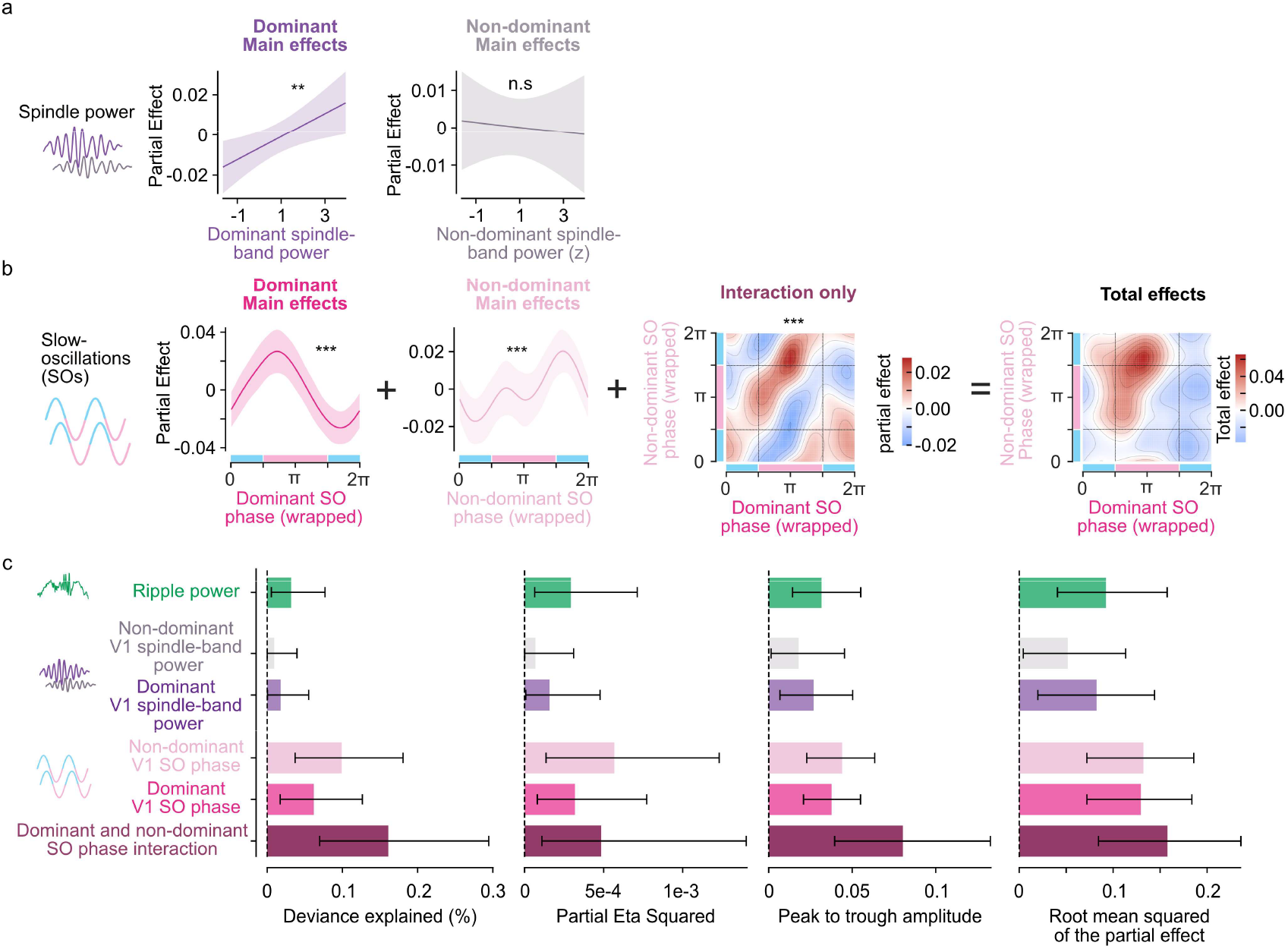
Contribution of each oscillatory predictor using full Generalized Additive Mixed Model (GAMM) including non-dominant hemisphere spindle-band power. An extended Generalized Additive Mixed Model (GAMM) was constructed including hippocampal ripple power, dominant- and non-dominant-hemisphere V1 spindle-band power, and dominant- and non-dominant-hemisphere V1 slow-oscillation (SO) phase simultaneously as predictors of per-event V1-HC reactivation coherence. **a**. Partial effects of V1 spindle-band power in dominant (left) and non-dominant (right) hemisphere on reactivation coherence. Shaded regions indicate 95% confidence intervals. **b**. Decomposition of the SO phase contribution into its constituent model terms: The main effects of SO phase in the dominant (first panel) and non-dominant (second panel) as well as the tensor-product interaction term (third panel) that explained the variance beyond the main effects. The total combined effect of all SO phase terms (fourth panel) is shown as a bivariate heatmap, with dominant SO phase on the x-axis and non-dominant SO phase on the y-axis. Phase windows corresponding to the SO peak and SO trough are indicated by green and pink shading on the respective axes. **c**. Four effect size metrics (deviance explained, partial *η*^2^, peak-to-trough amplitude, and RMS) for each predictor term, calculated based on 1000 case bootstrap replicates. Bars show the median bootstrapped RMS; error bars show the 95% bootstrapped confidence interval. Asterisks denote significance of each predictor in the model (\**p* < 0.05, \*\**p* < 0.01, \*\*\**p* < 0.001; ns, not significant).

**Extended Data Fig 8.**
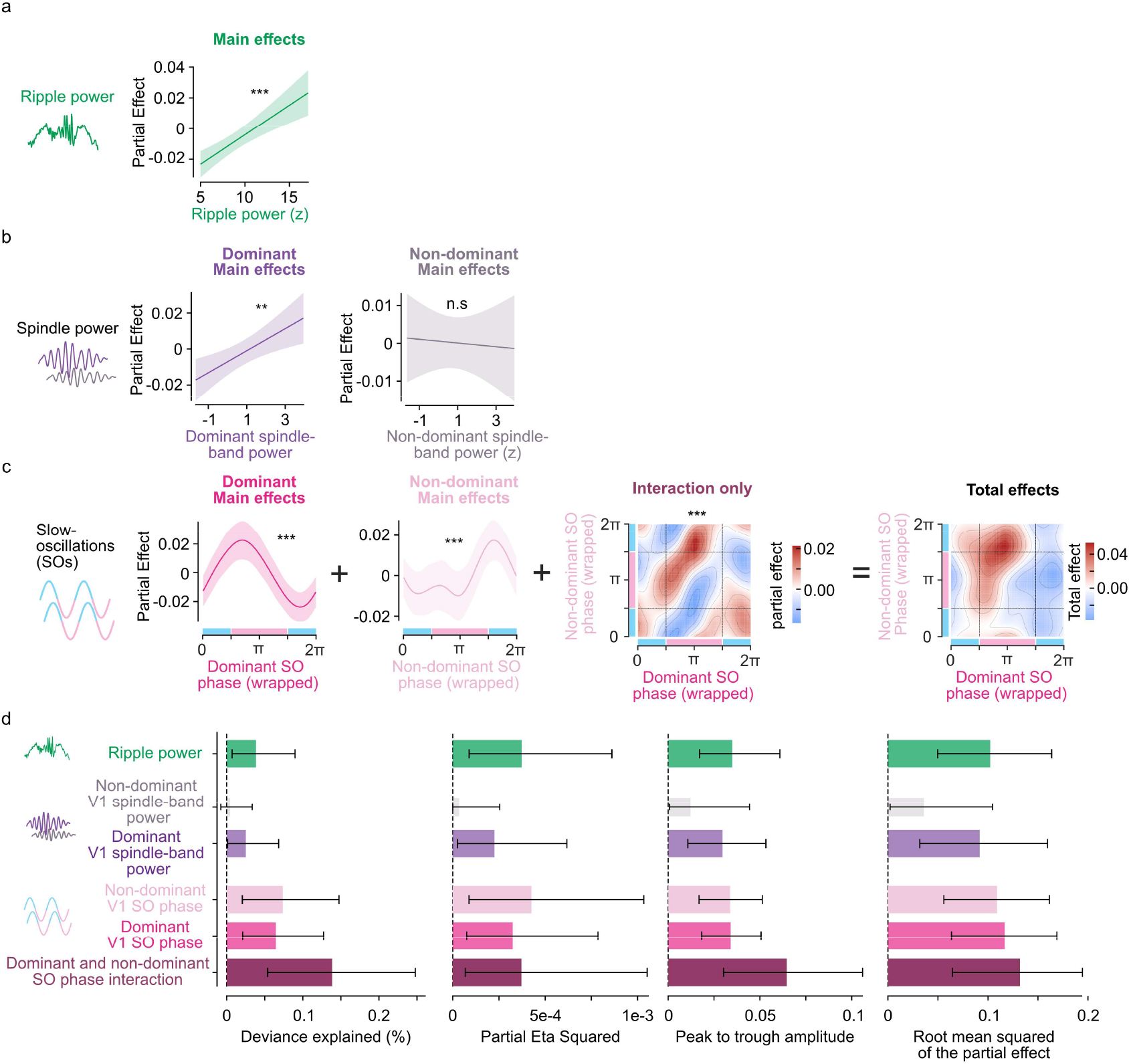
Generalized Additive Mixed Model (GAMM) results are robust to a wider window for quantifying reactivation coherence. The same model as Extended Data Fig 7 was applied. However, V1-HC reactivation coherence was calculated over a wider peri-ripple window of −200 to 200 ms, compared to the −0.1 to 0.1 s window used in Fig 5 and Extended data Fig 7.**a**. Partial effects of hippocampal ripple power on reactivation coherence. Shaded regions indicate 95% confidence intervals. **b**. Partial effects of V1 spindle-band power in dominant (left) and non-dominant (right) hemisphere on reactivation coherence. Shaded regions indicate 95% confidence intervals. **c**. Decomposition of the SO phase contribution into its constituent model terms: The main effects of SO phase in the dominant (first panel) and non-dominant (second panel) as well as the tensor-product interaction term (third panel) that explained the variance beyond the main effects. The total combined effect of all SO phase terms (fourth panel) is shown as a bivariate heatmap, with dominant SO phase on the x-axis and non-dominant SO phase on the y-axis. Phase windows corresponding to the SO peak and SO trough are indicated by green and pink shading on the respective axes. **d**. Four effect size metrics (deviance explained, partial *η*^2^, peak-to-trough amplitude, and RMS) for each predictor term, calculated based on 1000 case bootstrap replicates. Bars show the median bootstrapped RMS; error bars show the 95% bootstrapped confidence interval. Asterisks denote significance of each predictor in the model (\**p* < 0.05, \*\**p* < 0.01, \*\*\**p* < 0.001; ns, not significant).

## References

1. Buzsáki, G. Two-stage model of memory trace formation: A role for “noisy” brain states. Neuroscience 31, 551–570 (1989).

2. Frankland, P. W. & Bontempi, B. The organization of recent and remote memories. Nat Rev Neurosci 6, 119–30 (2005).

3. Diekelmann, S. & Born, J. The memory function of sleep. Nat Rev Neurosci 11, 114–126 (2010).

4. Wilson, M. A. & McNaughton, B. L. Reactivation of hippocampal ensemble memories during sleep. Science 265, 676–9 (1994).

5. Lee, A. K. & Wilson, M. A. Memory of Sequential Experience in the Hippocampus during Slow Wave Sleep. Neuron 36, 1183–1194 (2002).

6. Ji, D. & Wilson, M. A. Coordinated memory replay in the visual cortex and hippocampus during sleep. Nat Neurosci 10, 100–7 (2007).

7. Rothschild, G., Eban, E. & Frank, L. M. A cortical–hippocampal–cortical loop of information processing during memory consolidation. Nature Neuroscience 20, 251–259 (2017).

8. Sirota, A., Csicsvari, J., Buhl, D. & Buzsáki, G. Communication between neocortex and hippocampus during sleep in rodents. Proceedings of the National Academy of Sciences 100, 2065–2069 (2003).

9. Maingret, N., Girardeau, G., Todorova, R., Goutierre, M. & Zugaro, M. Hippocampo-cortical coupling mediates memory consolidation during sleep. Nat Neurosci 19, 959–64 (2016).

10. Staresina, B. P. et al. Hierarchical nesting of slow oscillations, spindles and ripples in the human hippocampus during sleep. Nat Neurosci 18, 1679–1686 (2015).

11. Leutgeb, S. et al. Independent Codes for Spatial and Episodic Memory in Hippocampal Neuronal Ensembles. Science 309, 619–623 (2005).

12. Steriade, M., Nunez, A. & Amzica, F. A novel slow (&< 1 Hz) oscillation of neocortical neurons in vivo: depolarizing and hyperpolarizing components. J. Neurosci. 13, 3252–3265 (1993).

13. Steriade, M., Domich, L. & Oakson, G. Reticularis thalami neurons revisited: activity changes during shifts in states of vigilance. J Neurosci 6, 68–81 (1986).

14. Buzsáki, G., Lai-Wo S. L. & Vanderwolf, C. H. Cellular bases of hippocampal EEG in the behaving rat. Brain Research Reviews 6, 139–171 (1983).

15. Latchoumane, C. V., Ngo, H. V., Born, J. & Shin, H. S. Thalamic Spindles Promote Memory Formation during Sleep through Triple Phase-Locking of Cortical, Thalamic, and Hippocampal Rhythms. Neuron 95, 424–435 e6 (2017).

16. Girardeau, G., Benchenane, K., Wiener, S. I., Buzsaki, G. & Zugaro, M. B. Selective suppression of hippocampal ripples impairs spatial memory. Nat Neurosci 12, 1222–3 (2009).

17. Robinson, H. L. et al. Large sharp-wave ripples promote hippo-campo-cortical memory reactivation and consolidation during sleep. Neuron 114, 226–236.e6 (2026).

18. Gridchyn, I., Schoenenberger, P., O’Neill, J. & Csicsvari, J. Assembly-Specific Disruption of Hippocampal Replay Leads to Selective Memory Deficit. Neuron 106, 291–300.e6 (2020).

19. Bendor, D. & Wilson, M. A. Biasing the content of hippocampal replay during sleep. Nat Neurosci 15, 1439–44 (2012).

20. Tirole, M., Huelin Gorriz, M., Takigawa, M., Kukovska, L. & Bendor, D. Experience-driven rate modulation is reinstated during hippocampal replay. eLife 11, (2022).

21. Steinmetz, N. A. et al. Neuropixels 2.0: A miniaturized high-density probe for stable, long-term brain recordings. Science 372, eabf4588 (2021).

22. Bimbard, C. et al. An adaptable, reusable, and light implant for chronic Neuropixels probes. eLife 13, (2025).

23. Huijeong Jeong, Vijay Mohan K. Namboodiri, Min Whan Jung, & Mark L. Andermann. Sensory cortical ensembles exhibit differential coupling to ripples in distinct hippocampal subregions. Current Biology https://doi.org/10.1101/2023.03.17.533028 (2023) doi:10.1101/2023.03.17.533028.

24. Siapas, A. G. & Wilson, M. A. Coordinated Interactions between Hippocampal Ripples and Cortical Spindles during Slow-Wave Sleep. Neuron 21, 1123–1128 (1998).

25. Xia, F. et al. Parvalbumin-positive interneurons mediate neocortical-hippocampal interactions that are necessary for memory consolidation. Elife 6, e27868 (2017).

26. Levenstein, D., Buzsáki, G. & Rinzel, J. NREM sleep in the rodent neo-cortex and hippocampus reflects excitable dynamics. Nat Commun 10, 2478 (2019).

27. Mölle, M., Yeshenko, O., Marshall, L., Sara, S. J. & Born, J. Hippocampal sharp wave-ripples linked to slow oscillations in rat slow-wave sleep. J Neurophysiol 96, 62–70 (2006).

28. Sirota, A. & Buzsáki, G. Interaction between neocortical and hippocampal networks via slow oscillations. Thalamus Relat Syst 3, 245–259 (2005).

29. Saleem, A. B., Chadderton, P., Apergis-Schoute, J., Harris, K. D. & Schultz, S. R. Methods for predicting cortical UP and DOWN states from the phase of deep layer local field potentials. J Comput Neurosci 29, 49–62 (2010).

30. Mohajerani, M. H., McVea, D. A., Fingas, M. & Murphy, T. H. Mirrored Bilateral Slow-Wave Cortical Activity within Local Circuits Revealed by Fast Bihemispheric Voltage-Sensitive Dye Imaging in Anesthetized and Awake Mice. J. Neurosci. 30, 3745–3751 (2010).

31. Vyazovskiy, V. V. et al. Local sleep in awake rats. Nature 472, 443–447 (2011).

32. Seibt, J. et al. Cortical dendritic activity correlates with spindle-rich oscillations during sleep in rodents. Nat Commun 8, 684 (2017).

33. Niethard, N., Ngo, H.-V. V., Ehrlich, I. & Born, J. Cortical circuit activity underlying sleep slow oscillations and spindles. Proceedings of the National Academy of Sciences 115, E9220–E9229 (2018).

34. Diamanti, E. M. et al. Spatial modulation of visual responses arises in cortex with active navigation. eLife 10, (2021).

35. Saleem, A. B., Diamanti, E. M., Fournier, J., Harris, K. D. & Carandini, M. Coherent encoding of subjective spatial position in visual cortex and hippocampus. Nature 562, 124–127 (2018).

36. Saleem, A. B. & Busse, L. Interactions between rodent visual and spatial systems during navigation. Nat Rev Neurosci 24, 487–501 (2023).

37. Morimoto, M. M., Uchishiba, E. & Saleem, A. B. Organization of feedback projections to mouse primary visual cortex. iScience 24, 102450 (2021).

38. Fournier, J. et al. Mouse Visual Cortex Is Modulated by Distance Traveled and by Theta Oscillations. Curr Biol 30, 3811–3817.e6 (2020).

39. Zutshi, I. & Buzsáki, G. Hippocampal sharp wave ripples and their spike assembly content are regulated by the medial entorhinal cortex. Curr Biol 33, 3648–3659.e4 (2023).

40. Lewis, P. A. & Bendor, D. How Targeted Memory Reactivation Promotes the Selective Strengthening of Memories in Sleep. Current Biology 29, R906–R912 (2019).

41. Meer, M. A. A. van der & Bendor, D. Awake replay: off the clock but on the job. Trends in Neurosciences 48, 257–267 (2025).

42. Massimini, M., Huber, R., Ferrarelli, F., Hill, S. & Tononi, G. The sleep slow oscillation as a traveling wave. J Neurosci 24, 6862–6870 (2004).

43. Nir, Y. et al. Regional Slow Waves and Spindles in Human Sleep. Neuron 70, 153–169 (2011).

44. Göldi, M., van Poppel, E. A. M., Rasch, B. & Schreiner, T. Increased neuronal signatures of targeted memory reactivation during slow-wave up states. Sci Rep 9, 2715 (2019).

45. Batterink, L. J., Creery, J. D. & Paller, K. A. Phase of Spontaneous Slow Oscillations during Sleep Influences Memory-Related Processing of Auditory Cues. J Neurosci 36, 1401–1409 (2016).

46. Cheng, S. & Frank, L. M. New experiences enhance coordinated neural activity in the hippocampus. Neuron 57, 303–313 (2008).

47. Huelin Gorriz, M., Takigawa, M. & Bendor, D. The role of experience in prioritizing hippocampal replay. Nat Commun 14, 8157 (2023).

48. Chang, H. et al. Sleep microstructure organizes memory replay. Nature 637, 1161–1169 (2025).

49. Singer, A. C. & Frank, L. M. Rewarded Outcomes Enhance Reactivation of Experience in the Hippocampus. Neuron 64, 910–921 (2009).

50. Lopes, G. et al. Creating and controlling visual environments using BonVision. eLife 10, e65541 (2021).

51. Lopes, G. et al. Bonsai: an event-based framework for processing and controlling data streams. Front Neuroinform 9, 7 (2015).

52. Buccino, A. P. et al. SpikeInterface, a unified framework for spike sorting. eLife 9, e61834 (2020).

53. Pachitariu, M., Sridhar, S., Pennington, J. & Stringer, C. Spike sorting with Kilosort4. Nat Methods 21, 914–921 (2024).

54. Fabre, J. M. J. Beest, E. H. van, Peters, A. J., Carandini, M. & Harris, K. D. Bombcell: automated curation and cell classification of spike-sorted electrophysiology data. Zenodo 10.5281/ze-nodo.8172822 (2023).

55. Pompili, M. N. & Todorova, R. Discriminating Sleep From Freezing With Cortical Spindle Oscillations. Front. Neural Circuits 16, (2022).

56. Tingley, D. & Buzsáki, G. Routing of Hippocampal Ripples to Subcortical Structures via the Lateral Septum. Neuron 105, 138–149.e5 (2020).

57. Hazon, O. et al. Noise correlations in neural ensemble activity limit the accuracy of hippocampal spatial representations. Nat Commun 13, 4276 (2022).

58. Krishnan, A., Williams, L. J., McIntosh, A. R. & Abdi, H. Partial Least Squares (PLS) methods for neuroimaging: A tutorial and review. NeuroImage 56, 455–475 (2011).

59. Carey, A. A., Tanaka, Y. & van der Meer, M. A. A. Reward revaluation biases hippocampal replay content away from the preferred outcome. Nat Neurosci 22, 1450–1459 (2019).

60. Takigawa, M., Huelin Gorriz, M., Tirole, M. & Bendor, D. Evaluating hippocampal replay without a ground truth. eLife 13, e85635 (2024).

